# All the diversity we cannot see: Using single-cell population genetics to reveal the reproductive mode of the snow alga *Chlainomonas* sp. (Chlorophyta)

**DOI:** 10.64898/2026.11.19.767852

**Authors:** Alexander Geragotelis, Solenn Stoeckel, Robin B. Kodner, Trinity L. Hamilton, Honu Pata, Stacy A. Krueger-Hadfield

**Affiliations:** Virginia Institute of Marine Science Eastern Shore Laboratory, Batten School of Coastal & Marine Sciences, William & Mary, Wachapreague, VA, USA; DECOD, L’Institut Agro, IFREMER, INRAE, Rennes, France; Department of Environmental Science, Western Washington University, Bellingham, WA, USA; Department of Plant and Microbial Biology and The Biotechnology Institute, University of Minnesota, Saint Paul, MN, USA; Department of Biology, University of Alabama at Birmingham, Birmingham, AL, USA

**Keywords:** Algae, *Chlainomonas* (snow alga), Ploidy, Reproductive mode, Single cell population genetics, Snow

## Abstract

Many organisms responsible for globally important processes, from carbon cycling to human disease, are microscopic eukaryotes. While their reproduction is essential for their success, across the tree of life, our understanding of evolutionary processes is limited due to the inherent challenges of studying microorganisms. Tools and predictions developed in multicellular and obligately sexual taxa are often not tractable in microbes, where population sizes can be large and extracting DNA from individuals is difficult. Even when sexual characteristics can be observed, population genetics is the only method available to characterize the reproductive mode. The relative rates of sexual versus clonal reproduction partition genetic diversity and are therefore integral to our understanding of evolutionary processes. We developed a single-cell population genetic approach to characterize the reproductive mode of the snow alga *Chlainomonas* sp. This Chlorophyte alga exhibits a complex life cycle, whereby bloom dynamics are directly observable in natural populations. We explored genetic diversity using single cells collected across several years of *Chlainomonas* sp. summer blooms. Dominant red cells in the bloom were diploid. Yet, all were unique genotypes, despite the prediction that clonal processes drive bloom development. We found high selfing rates, but selfing and clonality are often indistinguishable without understanding an organism’s natural history. *Chlainomonas* sp. has a haploid-diploid life cycle as hypothesized, engaging in both clonal and sexual reproduction, likely selfing. This study represents a first use of single cell population genetic methods to understand natural populations of partially clonal microeukaryotes.

## Introduction

The relative rates of sexual versus clonal reproduction determine the partitioning of genetic diversity, thereby structuring populations and ultimately influencing evolutionary trajectories through phenotypic evolution (Tibayrenc & Ayala, 1991; Hamrick & Godt, 1996; Orive *et al*., 2017). Yet, our understanding of evolutionary processes in partially clonal eukaryotes is limited due to the inherent challenges of studying these taxa using theoretical and analytical frameworks developed primarily for strictly sexual or strictly clonal organisms, making them difficult to apply to populations engaging in *both* sexual and clonal reproduction (Halkett *et al*., 2005; Arnaud-Haond *et al*., 2020; Stoeckel *et al*., 2021a). Partial clonality has implications for the behavior of commonly used population genetic summary statistics (Balloux *et al*., 2003; Arnaud-Haond *et al*., 2007; Stoeckel *et al*., 2021b; Krueger-Hadfield *et al*., 2021a). For example, excess heterozygosity is often used as evidence of clonal reproduction (Balloux *et al*., 2003). However, negative values of the inbreeding coefficient (*F_IS_* < 0; i.e., excess heterozygosity) can be found in sexual populations with no evidence of clonality (Becheler *et al*., 2017), and the variance in *F_IS_* is subject to the effects of sampling intensity (Arnaud-Haond *et al*., 2020; Stoeckel *et al*., 2021a; Stoeckel *et al*., 2021b). For life cycles with haploid phases in which mitoses occur (e.g., unicellular dinoflagellates; many macroalgae with macroscopic and independent haploid phases), *F_IS_* is meaningless and therefore, this commonly reported statistic in the literature is not available for comparison to other taxa (see as an example, Olsen *et al*., 2020). Moreover, the proportion of the haploid phase in haploid-diploid life cycles, such as found in mosses and many micro- and macroalgae, will have a disproportionate influence on patterns of intraspecific diversity (Stoeckel *et al*., 2021b). A small haploid proportion may increase the stochasticity of summary statistics used to characterize the reproductive mode (see Williams *et al*., 2026). Finally, signatures of selfing and clonality often converge because both reproductive modes constrain the effective recombination of genetic variation, thereby promoting linkage disequilibrium and the persistence of multilocus associations (Nordborg *et al*., 2000; Tibayrenc & Ayala, 2012; Taylor *et al*., 2015; Jullien *et al*., 2019). Thus, the convergence of genetic signatures generated by different reproductive modes, together with the limitations of existing theoretical frameworks, renders it challenging to reliably disentangle reproductive modes in natural populations, particularly in partially clonal taxa and organisms with haploid or haploid–diploid life cycles.

These limitations are compounded for unicellular eukaryotes – many of which are found across evolutionarily diverse lineages, including taxa responsible for globally important carbon cycling (e.g., dinoflagellates, diatoms, haptophytes, cryptomonads, green microalgae), human disease (e.g., apicomplexans [*Plasmodium*], fungi [*Candida*], eulgenozoans [*Leishmania*], diplomonads [*Giardia*]), and agricultural pests and diseases (e.g., *Phytophthora, Monilinia*). Population genetic studies have revealed the importance of clonal reproduction in structuring populations of some parasitic protozoa (e.g., Tibayrenc & Ayala, 1991; Tibayrenc, 1997), whereas other taxa, such as agricultural pests, engage in sexual reproduction on a cyclical basis that is interspersed with clonal processes (e.g., Everhart & Scherm, 2015). Often for these taxa, genotyping is performed following *in vitro* cultivation methods that can be subject to mutations, contamination risks, or the inability for a given genotype or species to survive in laboratory conditions. Consequently, population genetic data are missing for most microbial eukaryotes despite being the only way to describe the relative contribution of sexual versus clonal reproduction.

Additional considerations in microbial eukaryotes further complicate population genetic approaches. In microscopic eukaryotes, generation times are often short (Romeijn & Smallegange, 2024) and populations can be large (Gernstein & Moore, 2011). Previously, there has been a reliance on the isolation of single cells from the environment that are then subjected to clonal amplification under artificial laboratory conditions to obtain sufficient biomass for genotyping of ‘individuals’ in the unitary organism sense (e.g., 30 fish; see examples in haptophytes: [Krueger-Hadfield *et al*., 2014]; dinoflagellates: [Dia *et al*., 2014]; apicomplexans: [Ajzenberg *et al*., 2002]; phytophthorans: [Flier *et al*., 2003]). Thirty clonal cultures are hardly representative of populations that can comprise tens of thousands of cells per milliliter. Currently, only ∼1% of microbes can be cultured (Dawson & Pace, 2002; del Campo *et al*., 2014), thereby rendering it impossible to collect data for most microbial species. Moreover, during the establishment of cultures, many cultures fail to survive, and allelic shifts have been detected, raising questions about how representative of natural diversity these approaches are (see Krueger-Hadfield *et al.,* 2014 and references therein). Thus, bypassing this *in vitro* clonal amplification step is a major methodological objective for characterizing reproductive modes and thereby enabling the empirical observation of evolution in natural populations of microbial eukaryotes.

Recent technological advances have led to commercially available whole genome amplification (WGA) kits that enable genomic DNA amplification from single cells, thereby improving genome assemblies in apicomplexans (Morris *et al*., 2019) and trypanosomatids (Imamura *et al.,* 2020). WGA can also support single cell population genetics, as it allows for genotyping a single cell and therefore meeting assumptions of population genetic models (i.e., individuals). This would allow population studies to assess the population genetics of unicellular taxa, especially for the unculturable majority, where cells can be isolated from field samples. Thus, the population genetics of microbial taxa can proceed in a similar manner to macroscopic taxa and reveal the evolutionary ecology of these organisms – a major goal for the community of microbial biologists. Gollnisch *et al.,* (2024) used WGA to assess intraspecific population diversity and found unique genotypes from cells isolated from wild blooms, providing insight into the genetic diversity of the freshwater microalga *Gonyostomum semen* in both its native and non-native range.

While single nucleotide polymorphisms (SNPs) have become more common for population genetic studies, reduced representation approaches require knowledge about the ploidy of each specimen to assign a heterozygous or homozygous allelic state (e.g., Gompert *et al.,* 2014). For many taxa, ploidy is not known because phases are indistinguishable, such as in macroalgae, or because the life cycle is not known, such as in many microbes. These approaches often prioritize sample size within or among populations over sequencing depth, consequently relying on genotype likelihoods (e.g., Gompert *et al.,* 2014). Moreover, not all samples will be genotyped at the same loci using common approaches, such as restriction-site associated (RAD-Seq) genotyping by sequencing (GBS). For example, Flanagan *et al*. (2021) relied on microsatellite loci to establish ploidy and assign clonal lineage in the red macroalga *Gracilaria vermiculophylla* prior to performing analyses of population structure. These GBS approaches are applicable to sexual organisms whereby alleles are constantly shuffled each generation. In contrast, in partially clonal taxa, such as *G. vermiculophylla* (Krueger-Hadfield *et al*., 2016; Oetterer *et al*., 2026), the entire multilocus genotype can be passed on via clonal reproduction. Thus, typical SNP genotyping does not easily transfer to partially clonal taxa in which each specimen requires genotyping at the same suite of loci to characterize clonal lineages. In contrast, microsatellite genotyping, or SNPs using specifically targeted sites approaches such as HiPlex (Besnard *et al*., 2023), allow for the genotyping of every locus for each specimen, and therefore more rigorous assignment of clonal lineages. Combined with advances in WGA of single cells, studying the population genetics of unicellular taxa is now tractable from locus development to population-level analyses.

We have leveraged WGA to explore the reproductive mode of *Chlainomonas* sp. from the Cascades Mountains (Chlorophyta, Chlamydomonadaceae) – a snow algal taxon with predictable seasonal blooms composed solely *Chlainomonas* sp. rendering it amenable for population studies. *Chlainomonas* is one of the major genera of snow algal species inhabiting seasonal snowpack, blooming in spring and summer months when there is sufficient water available in the snow in alpine environments (Hoham, 1974a; Hoham, 1974b; van Hees *et al*., 2023). Snow and glacier ice algae are adapted to dynamic seasonal and annual changes in available habitat and are known to play notable roles in the intensity and duration of the melt season by reducing the surface albedo and are facing dramatic changes to habitat as the cryosphere warms (Williamson *et al*., 2020; Roussel *et al*., 2024). However, as these alpine habitats are undergoing rapid change with annual snowpack decreasing and the melt season lengthening (Roop *et al*., 2020), understanding the role of these algae in snowmelt is of greater importance for forecasting future changes (Roussel *et al*., 2024). It is critical to understand reproduction in these organisms, but difficulties in culturing these organisms and the remote location of blooms make fine-scale population surveys challenging.

*Chlainomonas* sp. forms annual blooms in the ephemeral snow-on-lake habitat at Bagley Lake, near Mt. Baker in the Northern Cascades, USA (Figure 1, Table S1). Matsumoto *et al.,* (2024) proposed its life cycle based on extensive field and lab observations, wherein the annual summer blooms are made of large red, putatively diploid cells that undergo mitotic divisions at the snow surface. A unique form of external cell division has previously been observed in these large red cells (Matsumoto *et al*., 2024). Once the snow melts, cells sink to lake sediments where it is hypothesized that meiosis results in small red-green pill cells that are putatively haploid and survive the winter in lake sediments. The following spring, the pill cells undergo fertilization to generate new, flagellated red cells either on the snow surface, or that can swim up through slush to the snow surface and begin the new season’s blooms. The annual *Chlainomonas* sp. blooms at Bagley Lake likely consist of the single, undescribed *Chlainomonas* sp. (see van Hees *et al*., 2023; Novis *et al*., 2023) and recur seasonally from approximately May to July (Matsumoto *et al*., 2024). The visible, recurrent blooms of a single species provide an advantage of using *Chlainomonas* sp. to develop WGA techniques and facilitate repeated sampling of the same putative population without the risk of sampling more than one microalgal species.

**Figure 1.**
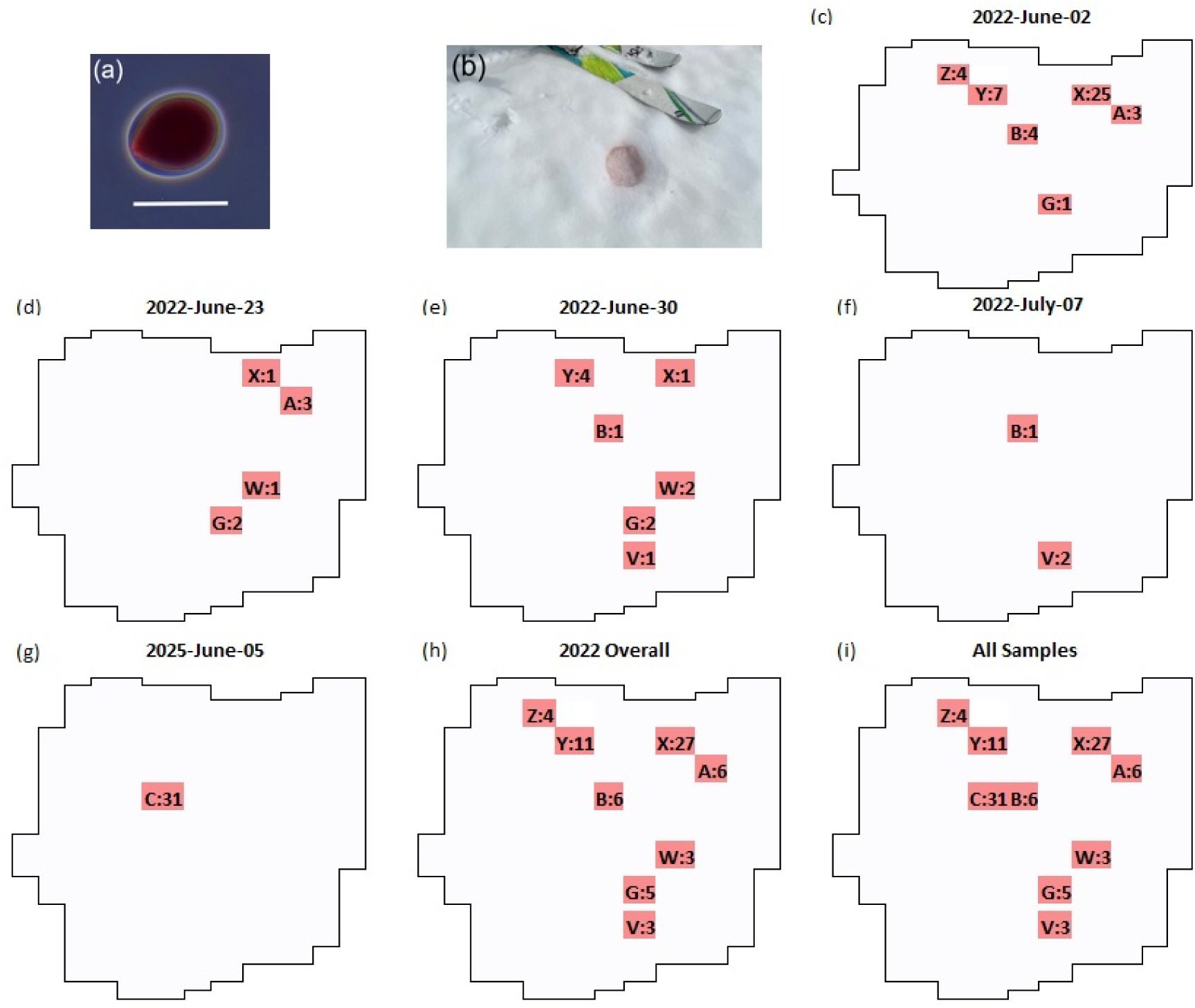
(a) An individual *Chlainomonas* sp. cell representative of cells forming blooms (scale bar = 50 μm) and amplified using PicoPLEX. (b) In front of the ski, there is a patch formed by a *Chlainomonas* sp. bloom, forming a cuvette in the snow (scale bar = 10 cm). (c) Map showing the relative locations of sampling areas on Upper Bagley Lake sampled on 2 June 2022. The letter-number combination shows the site name and the number of cells successfully genotyped and included in the final dataset. (d-i) Map showing the relative locations of sampling areas on Upper Bagley Lake sampled (d) on 23 June 2022; (e) on 30 June 2022; (f) on 7 July 2022; (g) on 5 June 2025; (h) throughout 2022; (i) in both 2022 and 2025. Photo credit (a-b): R. Kodner.

Here, we explored population diversity and reproduction in this system using single-cell WGA methods. As previous efforts to determine *Chlainomonas* sp. ploidy via DAPI staining were unsuccessful (Matsumoto *et al*., 2024), we first tested the hypothesis that the large red cells in the visible bloom were diploid. Chlorophycean lineages, including snow algal taxa (but see Hoham & Remias, 2020), have long been thought to have haploid life cycles, in which mitosis occurs solely in the haploid phase (Beukeboom & Perrin, 2014; Arora & Sahoo, 2015). While life cycle diversity in green algae has long been suspected (Otto & Marks, 1996), concrete evidence has only recently emerged. For example, Li *et al.,* (2026) provided genomic evidence for haploid-diploid life cycles in the Scenedesmaceae, suggesting ample life cycle variation within this unicellular lineage of Chlorophyceae as found in the Ulvophyceae (Otto & Marks, 1996). Then, we interpreted population genetic indices and Bayesian estimates of quantitative reproductive modes using changes in genotype frequencies (Stoeckel *et al*., 2024a) to explore the prediction that the population of *Chlainomonas* sp. at Bagley Lake is partially clonal. Both diploid and haploid cells are thought to undergo mitotic divisions, and bloom development likely follows similar clonal expansion as found in other microalgae (e.g., diatoms: [d’Alelio *et al*., 2010]; haptophytes: [Krueger-Hadfield *et al*., 2014]). This study is an important test of WGA for downstream population genetic analyses of individual cells isolated from the environment (see Gollnisch *et al*., 2024) as well as increasing our understanding of snow algal reproductive modes. While observations such as those provided in Matsumoto *et al.,* (2024) or reviewed in Hoham & Remias, (2020) provide necessary natural history data as well as hypothesis generation, this study begins to address details of reproduction that are necessary to understand the evolution in this algal group. These methods can be used on any uncultivatable microbial taxa and show promise to reveal many of the hidden facets of microbial life, which are often taken for granted in other taxa (e.g., animals or some angiosperms).

## Materials and Methods

### Sample collection, cell sorting, and DNA amplification

We sampled *Chlainomonas* sp. patches around Upper Bagley Lake near Mt. Baker in the North Cascades (48°51’14.8"N 121°41’30.5"W) using the grid established by van Hees *et al*., (2023) from the 2021 field campaign. The bloom in 2022 was less intense than the bloom in 2021, and the sampling area had to be expanded to retrieve more biomass (Figure 1). Each patch was sampled within a 1 m diameter around a marker designating an area of the lake (A, B, G, W, V, X, Y, Z for 2022 and C for 2025; Figure 1, Table S1). If obvious biomass was observed (i.e., pink snow), each patch was sampled weekly. Once a patch had no observable biomass, a new patch in the same sampling area was established. A 50 mL falcon tube was used to scrape along the snow surface, collecting visibly pink snow at each patch.

Single *Chlainomonas* sp. cells were isolated manually using an Olympus SZ61 dissecting scope (Olympus Corporation, Tokyo, Japan) or a Nikon SMZ-745T Trinocular Digital HD Stereo Microscope 0.67x - 5.0x (Nikon Corporation, Tokyo, Japan) by pipette and directly placed into a single well of a 96-well plate. After sorting, plates were stored at −20°C until whole genome amplification.

Genomic DNA was amplified using the PicoPLEX Single Cell Whole Genome Amplification (WGA) Kit (Takara Holdings, Kyoto, Japan) following version 3 of the manufacturer’s protocol. WGA DNA was diluted 1:100 with molecular grade water (Invitrogen, Waltham Massachusetts, USA).

### Microsatellite development, PCR, and fragment analysis

SSR-enriched genomic sequence data were generated by Microsynth Ecogenics GmbH (Balgach, Switzerland). We identified putative loci from the SSR-enriched library and followed Schoebel *et al*., (2013), with modifications implemented in Ryan *et al*., (2021) and Heiser *et al*., (2023). We used MSATCOMMANDER 1.0.8-beta (Faircloth, 2008) to design primers for di-, tri- and tetranucleotide repeat motifs separately. A minimum of eight repeats was selected and the following primer melting temperatures (T_m_): minimum of 58°C, optimum of 60°C, and maximum of 62°C. For dinucleotides, we identified 48,139 sequences with eight or more repeats; 9,608 of those had primers assigned, and 4,581 were potentially duplicated in the library. For trinucleotides, we identified 796 sequences with eight or more repeats, 326 of those had primers assigned, and 57 were potentially duplicated in the library. For tetranucleotides, we identified 11,782 sequences with six or more repeats, 1,259 of those had primers assigned, and 365 were potentially duplicated in the library. We had 5,027, 269, and 894 potential loci with di-, tri-, and tetranucleotide repeat motifs, respectively.

We used the R code provided by Schoebel *et al*., (2013) in R version 4.2.1 (R Core Team 2022) to combine the primer and microsatellite sequences into one file. For the dinucleotides, after merging the files we had 3,966 unique reads left. After removing duplicated forward and reverse primer sequences, we had 3,105 unique reads left. For trinucleotides, after merging the files we had 282 unique reads left. After removing duplicated forward and reverse primer sequences, we had 267 unique reads left. For tetranucleotides, after merging the files we had 828 unique reads left. After removing duplicated forward and reverse primer sequences, we had 684 unique reads left. We then combined the files with unique reads.

We calculated the absolute difference between the forward and reverse T_m_ for each primer pair and sorted all sets with an absolute difference between the forward and reverse T_m_ less than 1. We then sorted the putative loci by the forward penalty score, reverse penalty score, and the pair penalty score. Lastly, we calculated and sorted by the absolute difference between penalty scores divided by the pair penalty from smallest to largest to ensure that the difference between the forward and reverse penalties was as small as possible. We chose the top loci from each of those four categories and combined them in one file. We ranked the resulting 1,119 loci through the combined score from all four categories. Finally, we conducted a BLAST search of the first 100 of the primer pairs in Geneious Prime v.2022.2.2 (Biomatters, Ltd., Auckland, New Zealand) using the SSR-enriched library to ensure that only one primer pair was binding to the same locus, no primer pair was binding to more than one locus, and repeat regions were not within the primers.

We used eight *Chlainomonas* sp. cells sampled on 5 June 2025 from which WGA DNA was diluted 1:100 (Table S2). We performed PCR using either a SimpliAmp™ Thermal Cycler (Applied Biosystems, Waltham Massachusetts, USA) or a ProFlex PCR System (Applied Biosystems, Waltham Massachusetts, USA). PCR mix was prepared and a final volume of 15μL: 2μL template DNA, 250nM forward primer, 250nM reverse primer, 1X DreamTaq DNA Polymerase PCR Buffer with 20mM MgCl_2_ (Invitrogen Cat No. 18067017), 250nM of each dNTP (Thermo Scientific Cat No. R0192), 1 U DreamTaq DNA Polymerase (Thermo Scientific Cat No. EP0701). The following PCR program was used: 95 °C for 2 minutes followed by 35 cycles of 95 °C for 30 seconds, 58°C for 30 seconds, and 72 °C for 30 seconds, followed by a final extension of 72 °C for 5 minutes. PCR products were visualized by gel electrophoresis in a 1.5% agarose gel stained with GelRed (Millipore Sigma Cat No. SCT123). Only loci with a single band at the expected size range were retained for further testing. There were 16 putative loci that did not amplify and were eliminated. There were 14 loci that were eliminated due to having multiple bands when visualized on gel electrophoresis.

For fragment analysis, the PCR contained the same concentrations as shown above except for the labeled forward primer that had a fluorescent dye (6-FAM, NED, or PET, Table S3, S4, S5). PCR products were diluted in 9.7 µL of Hi-Di Formamide (Applied Biosystems, Cat No. 43-113-20) and 0.35 µL of the size standard GeneScan 500 LIZ (Applied Biosystems Cat No. 4322682) and fragment analysis was performed on a SeqStudio™ 24 Flex Sequencer (Applied Biosystems) at the Virginia Institute of Marine Science Eastern Shore Laboratory. Traces were visualized using Geneious Prime v.2025.1.2 (Biomatters, Ltd., Auckland, New Zealand). Bins for alleles at each locus were established based on manual allele calling (Table S4).

Seven loci were eliminated due to non-amplification following fragment analysis. There were a further 18 loci eliminated due to stutter peaks that rendered confident scoring difficult and may be indicative of amplifying multiple regions of the genome. The remaining twelve loci were then used for fragment analysis on the *Chlainomonas* sp. cells. Four of these loci were removed because they did not consistently amplify across cells from different sampling dates. Locus Ch28 was excluded as it was monomorphic. The seven final loci used in this study were polymorphic with 4 to 27 alleles (Table S4, S5). These seven loci were used to determine unique genotypes, as well as the proportion of unique genotypes in the population (Figure S1).

To determine the frequency of null alleles, we used the maximum likelihood estimator as implemented in ML-NullFreq (Kalinowski & Taper, 2006). The null allele proportions varied from 0 to 0.78 per locus per sampling date (Table S6). We note that other studies have found these estimates to be biased when populations are not mating at random (see Krueger-Hadfield *et al*., 2013).

### Single cell sample sizes

There were 267 individual *Chlainomonas* sp. cells amplified with PicoPLEX. Fifty-four cells (20.2%) were removed because they did not amplify at a single locus. There were 117 cells (43.8%) which amplified at four or fewer loci and were subsequently removed. Finally, there were 96 cells (36.0%) which amplified at five or more of the loci and were used to analyze the genetic structure in the Bagley Lake *Chlainomonas* sp. blooms in 2022 and 2025. These 96 cells were from 32 different patches across five sampling dates (Table S1).

### Data analyses

We calculated summary statistics using GenAPoPop 1.0 (Stoeckel *et al*., 2024a). The unbiased probability of identity (*pidu*) and the probability of identity between siblings (*pids*) were calculated to determine the resolution of the suite of microsatellite loci (Waits *et al.,* 2010). The probability of identity refers to the likelihood that two random individuals from a population will have the same genotype. A low value indicates that repeated genotypes are unlikely to arise in the population due to sexual reproduction, and consequently any repeated multilocus genotypes observed would likely have arisen from clonal reproduction.

Genotypic richness (*R*) was calculated as 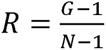, where G is the number of distinct genotypes and N is the total number of genotyped individuals (Dorken & Eckert, 2001). Genotypic evenness (*D\**) describes the distribution of sampling across clonal lineages (Ellstrand & Roose, 1987). Genotypic evenness is calculated as 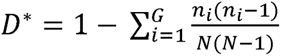, where *G* is the number of multilocus genotypes detected in the sample, and *n_i_* is the number of sampled units with the multilocus genotype (Arnaud-Haond *et al*., 2007). If a patch was dominated by few clonal lineages, *D\** approaches 0, and if every genet in the sampled area is represented by an equal number of ramets, *D\** approaches 1. Linkage disequilibrium (*r*□*_D_*) is a multilocus measure of the association of association of alleles (Agapow & Burt, 2001). Linkage disequilibrium is calculated with the formula 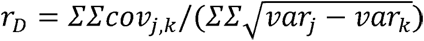, where *j* and *k* are two different loci. High *r*□*_D_* values indicate a lack of recombination, which can be a sign of clonal reproduction (Maynard Smith *et al.,* 1993). We calculated pairwise linkage disequilibrium (*LD*) using Hill multiallelic r^2^. This is calculated as 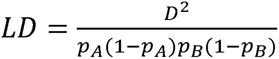, which normalizes linkage disequilibrium *D^2^,* to compare linkage disequilibrium between alleles of different frequencies. Next, we calculated the distribution of clonal membership (*Pareto β*) following Arnaud-Haond *et al.,* (2007). Based on available data in haploid-diploid mosses and macroalgae, Krueger-Hadfield et al. (2021a) denoted several thresholds: *Pareto β* values < 0.7 suggest high rates of clonal reproduction, values between 0.7 and 2.0 suggest partially clonal reproduction, and values > 2.0 suggest sexual reproduction. Allelic richness (*A_E_*) is a measure of the mean number of alleles observed at each locus (Weir, 1996). Allelic richness is calculated as *A_E_* 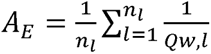, where *l* is the considered locus, *n_l_* is the number of the locus or marker used to produce the genotyping, and *Q_w,l_* is the population probability that two homologous alleles taken from different individuals in the population are identical. We calculated expected heterozygosity (*H_E_*), observed heterozygosity (*H_O_*), and the inbreeding coefficient (*F_IS_*) along with the variation in each index following Stoeckel *et al*., (2024b). Single locus *H_E_*, *H_O_*, and *F_IS_* values were calculated for the seven loci used to assess genotypes. We also calculated genetic differentiation (*F_ST_*) across time points. Finally, we calculated the relative rates of clonality, selfing, and outcrossing that could result in the observed genotype frequencies as implemented in GenAPoPop. We used the uniform joint priors covering the mutation rate u ∈ [0.001], the clonal rate *c* ∈ [0.0, 0.1, 0.2, 0.3, 0.4, 0.5, 0.6, 0.7, 0.8, 0.9, 1.0], and the selfing rate *s* ∈ [0.0, 0.1, 0.2, 0.3, 0.4, 0.5, 0.6, 0.7, 0.8, 0.9, 1.0] to cover the full spectrum of all possible quantitative reproductive modes. We considered outcrossing rates as the complement to one out of the clonal and selfing rates (0 = 1 – *c - s*).

For *r*□*_D_*, *A_E_*, *s*, *H_E_*, *H_O_*, and *F_IS_*, we performed linear regressions to assess temporal trends across sampling dates in R ver. 2026.05.0.

Figures were generated in R ver. 2026.05.0 using the following R packages ggplot2 (ver. 4.0.3; [Wickham, 2016]), ggimage (ver. 0.3.5; [Yu *et al*., 2017]), dplyr (ver. 1.2.1; [Wickham *et al*., 2026]), ggtext (ver. 0.1.2; [Wilke & Wiernik, 2002]), gridExtra (ver. 2.3; [Auguie, 2017]), cowplot (ver. 1.2.0; [Wilke, 2025]), patchwork (ver. 1.3.2; [Pedersen, 2025]).

### Microalgal *Pareto β* comparison

We performed a literature search on microalgae and population structure using Google Scholar. We focused on co-dominant markers: microsatellite loci and single nucleotide polymorphisms (SNP). We searched the key terms “microalgal population genetics microsatellite” and “microalgal population genetics snp”. We gathered 15 datasets, including five diatoms, seven dinoflagellates, one haptophyte, and two raphidophytes (Table S7). Seven of these datasets were publicly available, and the rest were provided by the corresponding authors. *Pareto β* was calculated for each population in each species using GenAPoPop 1.0 (Stoeckel *et al*., 2024a).

## Results and Discussion

We used whole genome amplification (WGA) for 267 *Chlainomonas* sp. cells and successfully genotyped 96 (36% success rate). Our results are comparable to Gollnisch *et al*., (2024). Previous microalgal studies have relied on clonal cultures initiated from single cells to generate sufficient biomass for DNA extraction and subsequent population genetic analyses (e.g., Iglesias-Rodríguez *et al*., 2006; Dia *et al*., 2014; Ruggiero *et al*., 2018). These analyses use bulk DNA and treat clonal cultures are an ‘individual’, thereby violating population genetic assumptions and thus, subsequent genotyping and interpretation are inherently limited. Here, we have demonstrated that single cells can be isolated from nature and directly genotyped as ‘individuals’ following WGA, adhering to one of the core assumptions of genetic theory – an individual (at least for genotyping even if that specimen is not a genet). However, as also demonstrated by Gollnisch *et al*., (2024), we experienced sizeable loss of *Chlainomonas* sp. cells following both WGA and PCR. Single-cell studies necessitate sampling substantially more cells than required for downstream analyses if ∼70% loss of isolated cells is common. While greater than losses associated with macroscopic algae (e.g., <10% of thalli removed due to poor DNA template or PCR amplification, see Williams *et al*., 2026; Oetterer *et al*., 2026), the losses observed by Gollnisch *et al*., (2024) or in our study are stochastic, likely reflecting cell preservation quality or cell lysing efficiency. By contrast, the initiation of laboratory cultures represents the artificial selection of clonal lineages that are either well suited or adapt more easily to *in vitro* conditions. Likewise, laboratory cultivation often imposes selection on cells that do well in a ‘colony’ and may select against cells that perform better as single cells. Thus, we argue that these losses from single cell population genetics are not the same as attempts to grow single cells in culture. Single cell genotyping represents a substantial step forward for understanding microbial evolutionary ecology.

Our seven microsatellite loci displayed sufficient power to resolve ploidy and genotypes (*pidu* = 1.12×10^-6^, *pids* = 5.81×10^-3^; Table 1). In our study, we observed heterozygosity in 79 (82%) of the 96 successfully genotyped cells (Table S2). While we were unable to determine the morphology of each red cell for which we performed WGA, initial sampling targeted red cells at the snow surface. Heterozygous genotypes at one or more loci support the proposal by Matsumoto *et al*., (2024) – red cells are diploid. We did not genotype the red-green pill cells that are thought to be haploid and are the product of meiosis. Nevertheless, if the dominant blooming cells are diploid, this supports the hypothesis of a haploid-diploid life cycle, such as found in other unicellular algae, like haptophytes (Frada *et al*., 2008; see also Beukeboom & Perrin, 2014). In the future, we need to genotype cells from across the life cycle to demonstrate ploidy differences as well as the clonal processes proposed by Matsumoto *et al*., (2024).

**Table 1.** Summary statistics based on 96 genotyped *Chlainomonas* sp. single cells by sampling date. *N*, sample size of genotyped cells; *pidu*, unbiased probability of identity of sibs; *pids*, probability of identity of sibs; *G*, number of genotypes; *R*, genotypic richness; *D\**, genotypic evenness; *Pareto β*, the distribution of clonal membership; *r*□_*D*_, estimate of linkage disequilibrium across loci; *A_E_*, mean observed allelic richness; *H_E_*, estimate of expected heterozygosity across loci; var(*H_E_*), the variance across loci observed in expected heterozygosity; *H_O_*, estimate of observed heterozygosity across loci; var(*H_O_*), the variance across loci observed in observed heterozygosity; *F_IS_*, estimate of the inbreeding coefficient across loci; var(*F_IS_*) the variance across loci observed in the inbreeding coefficient.

| **Sampling date** | ***N*** | ***pidu*** | ***pids*** | ***G*** | ***R*** | ***D\**** | ***Pareto β*** | ***r□<sub>D</sub>*** | ***A<sub>E</sub>*** | ***H<sub>E</sub>*** | **var(*H<sub>E</sub>*)** | ***H<sub>O</sub>*** | **var(*H<sub>O</sub>*)** | ***F<sub>IS</sub>*** | **var(*F<sub>IS</sub>*)** |
|---|---|---|---|---|---|---|---|---|---|---|---|---|---|---|---|
| 2 June 2022 | 44 | 7.50x10-<sup>7</sup> | 6.65x10<sup>-3</sup> | 44 | 1 | 1 | 4.5 | -0.005 | 3.7 | 0.599 | 0.055 | 0.240 | 0.025 | 0.600 | 0.037 |
| 23 June 2022 | 7 | nan | 7.77x10<sup>-3</sup> | 7 | 1 | 1 | 2.0 | -0.016 | 3.0 | 0.595 | 0.033 | 0.122 | 0.026 | 0.794 | 0.067 |
| 30 Jun 2022 | 11 | 1.73x10<sup>-5</sup> | 1.80x10<sup>-2</sup> | 11 | 1 | 1 | 2.6 | -0.030 | 2.4 | 0.511 | 0.030 | 0.186 | 0.005 | 0.637 | 0.094 |
| 5 June 2025 | 31 | 5.80x10<sup>-7</sup> | 6.94x10<sup>-3</sup> | 31 | 1 | 1 | 4.0 | -0.010 | 3.4 | 0.597 | 0.046 | 0.228 | 0.046 | 0.618 | 0.057 |

For 17 (18%) of the 96 genotyped cells, we observed a single allele at each of the seven genotyped loci (Table S2). In macroalgae, a single allele has been interpreted as either evidence for the haploid phase in isomorphic alterations of generations, or as fixed homozygosity in the diploid phase (e.g., Krueger-Hadfield *et al*., 2013). The combination of genotyping the red cells, positive inbreeding coefficients (*F_IS_*) (Table 1; Table S8; Figure 2a), and the life cycle proposed by Matsumoto *et al*., (2024) strongly suggest these 17 cells are fixed homozygotes. In other haploid-diploid macroalgae, high rates of selfing (i.e., self-fertilization) are common and result in homozygous diploid thalli (see Krueger-Hadfield *et al*., 2021a; Heiser *et al*., 2023; Krueger-Hadfield *et al*., 2024). In the case of the red macroalga *Chondrus crispus*, there were fixed homozygous thalli for which the diploid phase was confirmed by both the presence of reproductive structures and a chemical test that detects polysaccharide differences in the cell wall in haploids versus diploids (Krueger-Hadfield *et al*., 2013). By contrast, in the red macroalgae *Plocamium* spp. (Heiser *et al*., 2023) and *Chondria tumulosa* (Williams *et al*., 2026), for thalli without reproductive structures at the time of collection, there were no complementary chemical or genetic assays to determine ploidy (e.g., sex-linked markers, [Guillemin *et al*., 2012]). Based on population genetic indices and the overwhelming abundance of the diploid phase across populations, the authors hypothesized that many of the *Plocamium* and *Chondria* thalli with a single allele at each locus were fixed homozygous diploids (Heiser *et al*., 2023; Williams *et al*., 2026). In the case of *Chondria*, while clonal reproduction was the dominant reproductive mode, the authors saw an increase in homozygosity over time (Williams *et al*., 2024; Williams *et al*., 2026). While this may seem counterintuitive, as clonal reproduction is thought to be associated with increased heterozygosity (Balloux *et al*., 2003), in haploid-diploid taxa, the haploid proportion has an immense influence on genetic diversity (Stoeckel *et al*., 2021b). In *Chondria*, the haploid proportion was extremely low, and selfing was likely common when fertilization did occur, thereby increasing homozygosity against a background of clonal expansion in the diploid phase (Williams *et al*., 2026). Thus, in *Chlainomonas* sp., we suspect red cells are diploid, and meiosis occurs following snowmelt to produce the haploid red-green pill cells in the lake sediments. We propose that pill cells released from the same sporangium (see Figure 2 in Matsumoto *et al*., 2024) may readily undergo fertilization, in a process akin to intergametophytic selfing in multicellular taxa (Klekowski, 1969), and like the selfing in red macroalgae (e.g., *Chondrus crispus*, [Krueger-Hadfield *et al*., 2015; 2024]). Nevertheless, the recalcitrant cell walls of *Chlainomonas* sp. pose an issue for determining ploidy with nuclear staining methods, but future work must not only expand the cell types that are genotyped but also use additional methods to quantify ploidy (e.g., flow cytometry and chromosome counting using fluorescence *in situ* hybridization).

**Figure 2.**
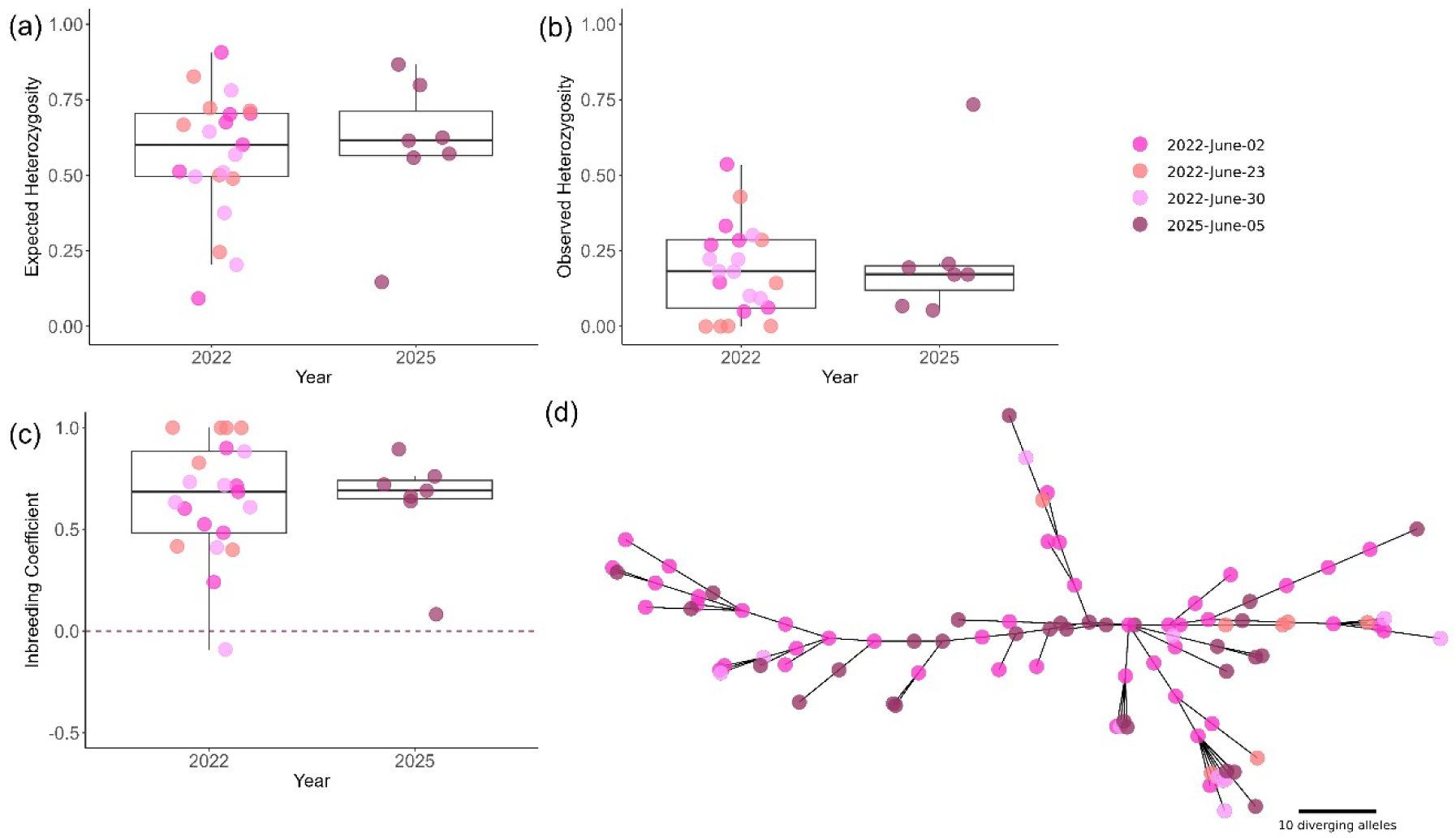
Genetic diversity in *Chlainomonas* sp. Boxplots by sampling date and year for (a) for single locus inbreeding coefficients (*F_IS_*). The red dashed line represents Hardy-Weinberg proportions that would be expected in a randomly mating population. (b) single locus estimates of expected heterozygosity (*H_E_*), (c) for single locus estimates of observed heterozygosity (*H_O_*). Boxes represent the interquartile range, the middle lines are medians, and whiskers represent the 1.5 interquartile ranges. Datapoints are jittered along the x-axes to improve visualization. (d) Minimum spanning tree for all sampling dates, illustrating the genetic relationships between individual cells. Points are colored in shades of pink by sampling date. Distance following connection between two individuals (points) is proportional to their shared ancestrality.

We expected clonal reproduction (i.e., mitotic division in red cells) to dominate the blooms in the *Chlainomonas* sp. population at Bagley Lake. Our suite of loci had sufficient power to resolve genotypes such that repeated genotypes would provide evidence for clonal processes during the seasonal blooms. Yet, every cell we genotyped was unique (Table S2). We sampled cells across 32 different patches across two years and often only had a single cell genotyped from a patch (Table S1). Notably, in patch C3, sampled in 2025, we successfully genotyped 31 cells from the same pink snow sample and all these cells were unique genotypes. We strongly suspect clonal processes via mitotic divisions of red cells facilitate and maintain the development of blooms turning the snow pink. Based on our results from patch C3, there is a vast diversity of clonal lineages in the Bagley Lake *Chlainomonas* sp. population. In other studies, sample sizes of 30 individuals have been recommended to describe genetic diversity (Ward & Jasieniuk, 2009). But even in a sample size of ∼30 cells, it is unlikely that we would sample repeated genotypes based on the large diversity of clonal lineages and when we have sampled so few cells relative to the total number of cells in a snow algal patch. Snow algal blooms contain up to thousands of cells per mL (Hoham & Remias, 2020). Therefore, it would be unlikely to find repeated genotypes when only a few cells were genotyped from an approximately 45 mL sample of pink snow. Our results also suggest repeated genotypes are not readily shared among patches. However, greater sampling intensity within and among patches is needed to verify if patches are unique clonal expansions. Nevertheless, our results, particularly from patch C3, help illustrate that the high rates of clonality in *Chlainomonas* sp. blooms do not necessarily mean there is a limited number of genomes, but rather that populations can be highly clonal while also containing a remarkable diversity of clonal lineages.

There have been proposals that microbial populations are ‘hyperdiverse’, containing significantly more genetic diversity than populations of macroscopic organisms, and consequently should be analyzed differently (Cutter *et al*., 2013). We argue that the requisite sample size to capture diversity and characterize clonal lineages in microbial taxa is logistically challenging, but this is not due to ‘hyperdiversity’. Hoban (2014) advocated for power analyses to calculate sufficient sample size to detect population structure, demonstrating for some taxa the sampling all individuals in a species would still be insufficient to characterize structure. When employed by Krueger-Hadfield & Hoban (2016) for haploid-diploid macroalgae, surprisingly large sample sizes were needed to capture sufficient sample sizes of haploid and diploid phases. This no doubt extends to uncovering repeated genotypes as well as microalgae. While some studies, such as in *Gephyrocapsa huxleyi* (as *Emiliania huxleyi*), have found repeated genotypes (Krueger-Hadfield *et al*., 2014), many microalgal studies have found unique genotypes for all samples (e.g., Dia *et al*., 2014; Gollnisch *et al*., 2024; Demura *et al*., 2014). In the case of the dinoflagellate *Alexandrium minutum* (Dia *et al*., 2014) or the diatom *Skeletonema marinoi* (Godhe & Härnström, 2010), high genetic diversity is likely generated by the resting cysts acting as a ‘seed bank’. These resting cysts allow for successful genotypes from previous years to simultaneously generate a bloom, increasing the observed genetic diversity within the bloom. It is possible that flagellated cells are swimming up to the snow surface from the sediments throughout the *Chlainomonas* sp. bloom. The staggered arrival of flagellated cells over the course of the summer bloom bears some similarities to *A. minutum* (Dia *et al*., 2014). Thus, the genetic diversity we observe in *Chlainomonas* sp. may be due to the genetic diversity accumulated in multiple years of sediment cysts in a manner like the ‘seed banks’ that generate *A. minutum* blooms. We do not know how long the red-green pill cells survive in lake sediments but based on our current results this is a fruitful avenue to pursue.

While not commonly applied in population genetic studies, we performed temporal analyses of genotype frequencies to quantify the relative rates of clonality, selfing, and outcrossing. We found high rates of selfing (*s*) between sampling dates (0.88 < *s* < 0.97), with low to zero rates of clonality and outcrossing (Table S9, S10). There was no evidence of a temporal trend in the selfing rate over the sampling period (linear regression: *β* = −2.20 × 10□□, 95% CI = −1.11 × 10 to 6.81 × 10□□, *p* = 0.540, *R*² = 0.100). While we suspect selfing occurs in lake sediments between pill cells originating from the same sporangium, it is unlikely that selfing dominates the population of *Chlainomonas* sp. at Bagley Lake, especially in the red cells we genotyped here. Further, if the *Chlainomonas* sp. population was primarily selfing, we would expect to see a decrease in expected heterozygosity (*H_E_*), observed heterozygosity (*H_O_*), and increasing inbreeding coefficient (*F_IS_*). Yet, we also observed no temporal trend in these three indices (linear regression: *H_E_β* = 2.53 × 10□□, 95% CI = −2.04 × 10□□ to 2.55 × 10□4, *p* = 0.682, *R*² = 0.101; *H_O_ β* = 4.06 × 10□□, 95% CI = −2.32 × 10□□ to 3.13 × 10□^4^, *p* = 0.588, *R*² = 0.170; *F_IS_ β* = −5.31 × 10□□, 95% CI = −5.27 × 10□□ to 4.21 × 10□^4^, *p* = 0.678, *R*² = 0.104; Table 1, Table S11; Figure 2a-c; Figure S2a-d). Further, stable linkage disequilibrium (*r□_D_*) would be expected in a clonal population, whereas selfing would result in changes in linkage disequilibrium as recombination would break associations among loci, with an overall higher *r□_D_* (Nordborg *et al*., 2000; de Meeûs *et al*., 2007; Stoeckel *et al*., 2024b). Similarly, no temporal trend in *r□_D_* was observed (linear regression: *β* = 6.09 × 10□^6^, 95% CI = −5.18 × 10□^5^ to 6.40 × 10□^5^, *p* = 0.695, *R*² = 0.093; Table 1, Figure S3). Selfing would also be expected to decrease allelic richness (*A_E_*) over time, as inbreeding erodes genetic diversity (Charlesworth & Charlesworth, 1987). However, during the 2022 bloom, there was no temporal pattern in allelic richness (linear regression: *β* = −0.045, 95% CI = −0.178 to 0.087, *p* = 144, *R*² = 0.950; Table 1). Finally, in the minimum spanning tree, we observed no clustering of 2025 genotypes compared to 2022 genotypes, despite the elapse of three years between sampling events (Figure 2d). These patterns might also reflect demographic differences between haploid and diploid phases that are not currently considered by current temporal-based methods (RMES for selfing [David *et al*., 2007] or ClonEstiMate [Becheler *et al*., 2017]). The lack of temporal trends in any of the population genetic indices, combined with stable genetic diversity, strongly suggests selfing does not exclusively dominate *Chlainomonas* sp. blooms.

While we expected high rates of clonality in *Chlainomonas* sp., the observation of unique genotypes and selfing from temporal analyses deviate from the expected effects of dominant clonality (Halkett *et al*., 2005). A single sexual event can reset genetic diversity but is unlikely to erase all signatures of clonality (Rouger *et al.,* 2016). Moreover, selfing and clonality can generate highly overlapping genetic signatures, making the two reproductive modes difficult to disentangle using existing population genetic approaches (e.g., Shainker-Connelly *et al*. 2024; Williams *et al*. 2026; see also Stoeckel *et al.,* 2024b). Thus, natural history observations become key to disentangling reproductive modes. Nevertheless, the patterns in our data for *Chlainomonas* sp. match other partially clonal microbial taxa. For example, the ascomycete fungus *Monilinia fructicola* affects stone fruits and clonal processes dominate their summer blooms, with signatures of significant linkage disequilibrium (Villarino *et al*., 2012). But when *M. fructicola* reproduces sexually, one bout of sex can yield many unique genotypes (Everhart & Scherm, 2015). The inocula at the start of each growing season are diverse and likely the result of both clonal and sexual reproduction (Everhart & Scherm, 2015). Similarly, in *Chlainomonas* sp., both temporal genetic differentiation (*F_ST_*) and pairwise locus linkage disequilibrium (*LD*) suggest the entry of new genotypes or one genetic group succeeding another over the course of the 2022 summer bloom. We found increasing genetic differentiation between 2 June and 30 June 2022, reaching two to eight times the genetic differentiation observed between two years in early June (Table S12). Similarly, pairwise *LD* decreased over the 2022 sampling dates, but not between 2022 and 2025 (Figure S4, Table S13). This is further supported by overall *LD* remaining lower than *LD* at each sampling date, indicating that there is more structure in *LD* on specific sampling dates than throughout the study period (Figure S4).

To further compare our hypotheses of the reproductive mode in *Chlainomonas* sp., we compared the distribution of clonal membership (*Pareto β*, [Arnaud-Haond *et al*., 2007]) across microalgal taxa. Based on empirical data from Krueger-Hadfield *et al*., (2021a) in macroalgae and mosses, *Pareto β* values < 0.7 indicate clonality, values between 0.7 and 2 indicate partial clonality, and values > 2 indicate sexual reproduction. In *Chlainomonas* sp., *Pareto β* values ranged from 2.0 to 4.5 across sampling dates (Table 1), not aligning with the expectations for a partially clonal species. However, many other partially clonal microalgae were found to have *Pareto β* values > 2 (Figure 3). At high rates of clonality in a homogeneous genetic group, the number of individuals of each genotype is expected to be skewed in a bimodal distribution. A smaller mode to the left corresponds to the new recombining lineages created by the rare sexual events and a larger mode to the right corresponds to the main clonal lineages growing over time. Together, this contributes to decreasing *Pareto β* values (which is the slope of this power-law inverse cumulative distribution). With the continuous entry of new genotypes or at least one phenological genetic group succeeding another over the course of the summer bloom as we suspect here in *Chlainomonas* sp., the small left mode of the size distribution of lineages would be artificially increased, while main lineages could not reach their expected sizes at a stationary distribution (Abdalrahem *et al*., 2025).

**Figure 3.**
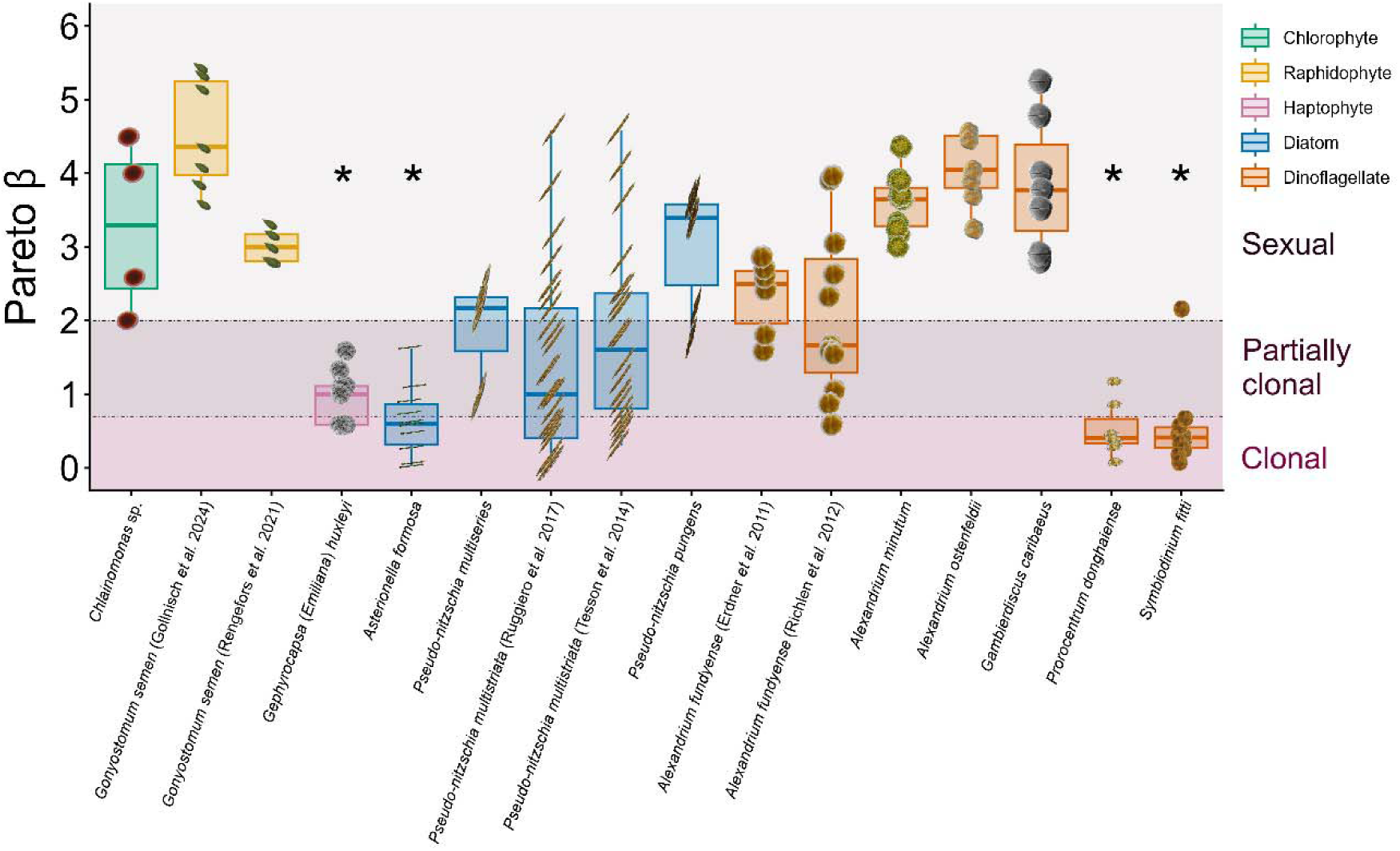
Boxplot of *Pareto *β** values calculated at different sampling dates in the 2022 and 2025 *Chlainomonas* sp. blooms compared to other microalgae organized by taxonomic group. Black asterisks indicate microalgae that do not form a ‘seed bank’ in which repositories of cells may be stored in sediments (e.g., *Alexandrium fundyense*) or on macroalgae (e.g., *Gambierdiscus caribaeus*). The dashed horizontal lines indicate the thresholds identified by Krueger-Hadfield et al. (2021a) as *Pareto β* ≤ 0.7 as clonal; 0.7 ≥ *Pareto β* ≤ 2 as partially clonal, and *Pareto β* ≥ 2 as sexual in haploid-diploid macroalgae and mosses. Data points are represented by images of each alga and are jittered along the x-axis to improve visualization. Data and image sources are provided in Table S7.

Classic population genetic theory has largely been developed for idealized populations characterized by diploidy and random mating in the form of sexual reproduction, as exemplified by the Wright–Fisher and standard coalescent frameworks (Fisher, 1922; Wright, 1931; Kingman, 1982). Although theoretical extensions have subsequently been developed to account for specific reproductive modes, such as selfing or clonality (Nordborg *et al*., 2000; Stoeckel & Masson, 2014; 2026), existing frameworks remain poorly suited to populations in which these reproductive modes coexist. We currently lack general mathematical models capable of predicting the joint genetic consequences of outcrossing, selfing, and clonality within diploid populations, and this limitation becomes even more pronounced in haploid–diploid life cycles, where these reproductive modes may operate in different phases of the life cycle. Recent work in freshwater (e.g., Shainker-Connelly *et al.,* 2024; Shainker-Connelly *et al*., 2025a; Shainker-Connelly *et al*., 2025b), polar (Heiser *et al*., 2023; 2025), and tropical red macroalgae (Williams *et al*., 2024; 2026) have all highlighted the difficulty of interpreting conventional population genetic statistics in the context of partial clonality and haploid-diploid life cycles. Our results here in *Chlainomonas* sp. as well as the observations of *Pareto β* in other microalgae further demonstrate that available analytical tools may fail to adequately characterize the prevailing reproductive mode in microalgae, and likely microbes in general. This limitation is particularly relevant for taxa such as *Chlainomonas* sp., which combine a haploid–diploid life cycle with the differentiation of mating types in the haploid phase (i.e., dioicy) and clonal reproduction in both haploid and diploid phases (see review in [Krueger-Hadfield, 2024]). Dioicy does not preclude selfing despite the presence of distinct sexes or mating types via intergametophytic selfing (Klekowski, 1969).

Limitations in both our understanding of the population genetics of haploid-diploid, partially clonal taxa and of the evolutionary ecology of microorganisms have previously been a barrier to research for uncultivatable microbial eukaryotes, such as snow algae. For this group of organisms, remote sensing studies (e.g., Lutz *et al*., 2016; Roussel *et al*., 2024) and phylogenetic research (e.g., Hoham & Remias, 2020; Procházková *et al.,* 2026) have elucidated parts of their biology and bloom dynamics. However, our inability to address basic questions about their life cycles and reproductive modes have limited our ability to forecast future changes in their populations (see Life Cycle section in Hoham & Remias, 2020). Our results show that novel techniques can allow us to address these questions, and further work can improve our understanding of *Chlainomonas* sp. and snow algae generally, as well as partially clonal eukaryotes more broadly. Partially clonal eukaryotes are found across evolutionarily diverse lineages, including taxa responsible for globally important processes. Our understanding of their evolution is essential to predicting the consequences of climate change in these systems.

This is also critical in organisms, such as *Chlainomonas* sp., that occupy habitats which are rapidly disappearing, with snow algal blooms possibly accelerating the loss of their own habitats (Cook *et al*., 2020; Williamson *et al*., 2020). For the unculturable majority of eukaryotes (Dawson & Pace, 2002; del Campo *et al*., 2014), single-cell genotyping promises a huge leap for understanding reproductive modes and life cycle variation, even for taxa for which culturing is possible, as maintaining full life cycles *in vitro* is challenging. The techniques here and in Gollnisch *et al.,* (2024) hold great potential to study evolution in response to environmental change. Using these population genetic tools to quantitatively assess the prevailing reproductive mode of populations will shed light on these species’ abilities to respond to environmental change through phenotypic evolution (Orive *et al*., 2017).

## Data availability

Data generated or analyzed during this study are included in this published article and its Supporting Information files. Microalgal *Pareto β* data in Figure 3 was generated from data is provided in the references.

## Author contributions

**ADG**: Data Curation (equal), Formal analysis (equal), Funding acquisition (supporting), Investigation (equal), Methodology (supporting), Visualization (equal), Writing – original draft (equal), Writing – review & editing (equal); **SS**: Formal analysis (equal), Investigation (equal), Methodology (supporting), Visualization (equal), Writing – original draft (supporting), Writing – review & editing (equal); **RBK**: Conceptualization (equal), Funding acquisition (equal), Investigation (supporting), Methodology (supporting), Resources (supporting), Supervision (supporting), Writing – review & editing (equal); **TLH**: Conceptualization (equal), Funding acquisition (equal), Methodology (supporting), Writing – review & editing (equal); **HP**: Investigation (supporting), Writing – review & editing (supporting); **SAKH**: Conceptualization (lead), Data curation (equal), Formal analysis (equal), Funding acquisition (lead), Investigation (equal), Methodology (lead), Resources (lead), Supervision (lead), Visualization (equal), Writing – original draft (equal), Writing – review & editing (equal).

## Acknowledgements & Funding

We are grateful to M. Matsumoto, C. Hanneman, A.G. Camara, S. Paradis, and D. van Hees for help with field work, as well as B. Thornton for initial microsatellite screening. We thank D. Erdner, M. Richlen, J. Jerney, R. Gollnisch, W-D. Yang, M. Montresor, M. Ruggiero, and N. Adams for supplying datasets of microalgal genotypes. We also thank A. Hardison, R. Carnegie, and J. McDowell for serving on the thesis committee for ADG. Finally, we thank the staff at the Virginia Institute of Marine Science Eastern Shore Laboratory for general support and infrastructure. This project was supported by the National Science Foundation (NSF) EAGER award DEB-2113745 to SAKH, DEB-2113746 to TLH, and DEB-2113747 to RBK, start-up funds from the College of Arts and Sciences at University of Alabama at Birmingham to SAKH, start-up funds from VIMS William & Mary to SAKH, and the Phycological Society of America Grant in Aid of Research Award to ADG. We also acknowledge support from the Owens Family Foundation Education Endowment for the Virginia Institute of Marine Science Eastern Shore Laboratory. Stipend support for ADG was provided by the Virginia Institute of Marine Science Office of Academic Affairs, the Juliette B. & Carroll W. Owens, Sr. Fellowship, and the James E. & Anne McCracken Rogers Graduate Student Fellowship.

