## Supplemental Information for "All the diversity we cannot see: Using single-cell population genetics to reveal the reproductive mode of the snow alga *Chlainomonas* sp. (Chlorophyta)"

The following Supporting Information is available for this article:

**Fig. S1** The proportion of unique genotypes identified in *Chlainomonas* sp. by polymorphic loci

**Fig. S2a** Violin plots of the number of raw alleles (*NA*) across sampling dates.

**Fig. S2b** Violin plots of the observed heterozygosity (*H_O_*) across sampling dates.

**Fig. S2c** Violin plots of the expected heterozygosity (*H_E_*) across sampling dates.

**Fig. S2d** Violin plots of the inbreeding coefficient (*F_IS_*) across sampling dates.

**Fig. S3** The trend in linkage disequilibrium (*r̄_D_*) values shown by each sampling date

**Fig. S4** Plot of pairwise linkage disequilibrium by locus using Hill multiallelic r^2^ across sampling dates

**Table. S1** Metadata and sample size for sampled patches of *Chlainomonas* sp.

**Table. S2** Microsatellite genotypes of *Chlainomonas* sp. cells.

**Table. S3** Primer development for Chlainomonas sp.

**Table. S4** Allele call table showing the locus, dye, raw allele range, and final allele call.

**Table. S5** Microsatellite locus information for *Chlainomonas* sp.

**Table. S6** Null allele proportions at each site and time point by locus.

**Table. S7** Studies used to compare population *Pareto β* to *Chlainomonas* sp*.*

**Table. S8a** Single locus values for expected heterozygosity (*H_E_*) at each patch and sampling date.

**Table. S8b** Single locus values for observed heterozygosity (*H_O_*) at each patch and sampling date.

**Table. S8c** Single locus values for the inbreeding coefficient (*F_IS_*) at each patch and sampling date.

**Table. S9a** Posterior probabilities of rates of clonality, selfing, and outcrossing using changes of genotype frequencies between temporal samples (CEMP) gathered from Bagley Lake on 2 June 2022 and 23 June 2022.

**Table. S9b** Posterior probabilities of rates of clonality, selfing, and outcrossing using changes of genotype frequencies between temporal samples (CEMP) gathered from Bagley Lake on 2 June 2022 and 30 June 2022.

**Table. S9c** Posterior probabilities of rates of clonality, selfing, and outcrossing using changes of genotype frequencies between temporal samples (CEMP) gathered from Bagley Lake on 2 June 2022 and 5 June 2025.

**Table. S9d** Posterior probabilities of rates of clonality, selfing, and outcrossing using changes of genotype frequencies between temporal samples (CEMP) gathered from Bagley Lake on 23 June 2022 and 30 June 2022.

**Table. S9e** Posterior probabilities of rates of clonality, selfing, and outcrossing using changes of genotype frequencies between temporal samples (CEMP) gathered from Bagley Lake on 23 June 2022 and 5 June 2025.

**Table. S9f** Posterior probabilities of rates of clonality, selfing, and outcrossing using changes of genotype frequencies between temporal samples (CEMP) gathered from Bagley Lake on 30 June 2022 and 5 June 2025.

**Table. S10** Intervals of inferred rates of clonality, selfing, and outcrossing from the genotype frequencies.

**Table. S11a** Significance of observed heterozygosity distribution between sampling dates.

**Table. S11b** Significance of expected heterozygosity distribution between sampling dates.

**Table. S11c** Significance of the inbreeding coefficient distribution between sampling dates.

**Table. S12** Temporal *F_ST_* between sampling dates.

**Table. S13** Significance of pairwise linkage distribution (LD) between sampling dates.

**Fig. S1**

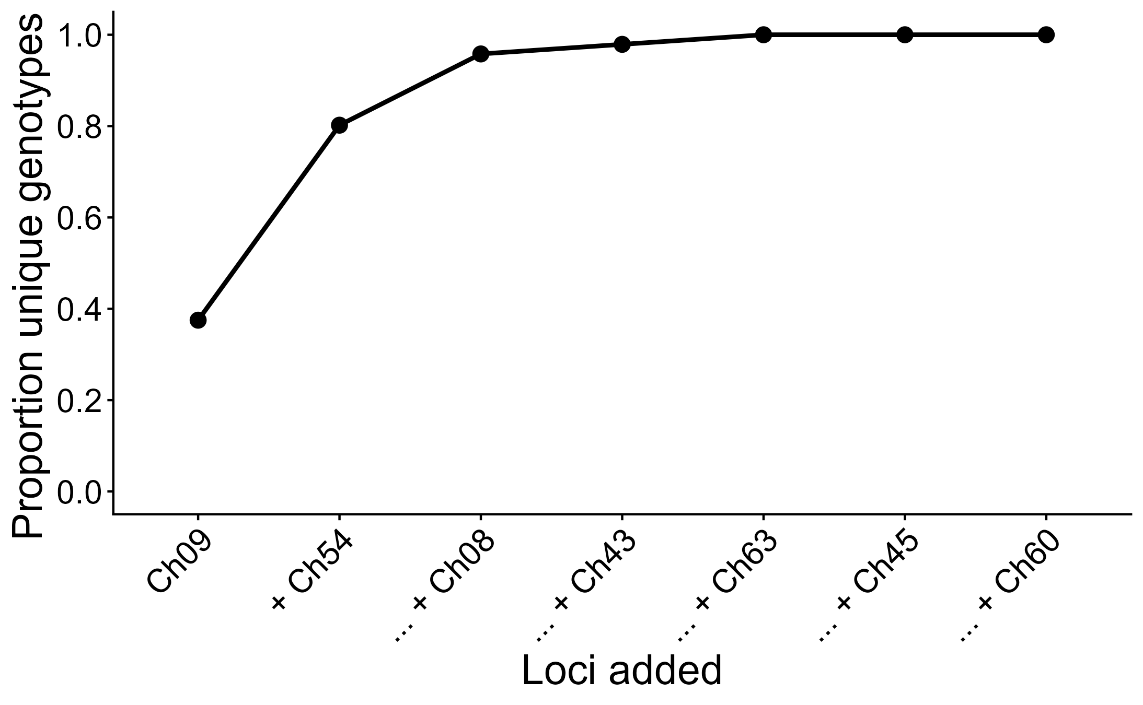

Figure S1. The proportion of unique genotypes identified in *Chlainomonas* sp. when adding microsatellite loci from the most polymorphic to least polymorphic based on allelic richness.

**Fig. S2a**

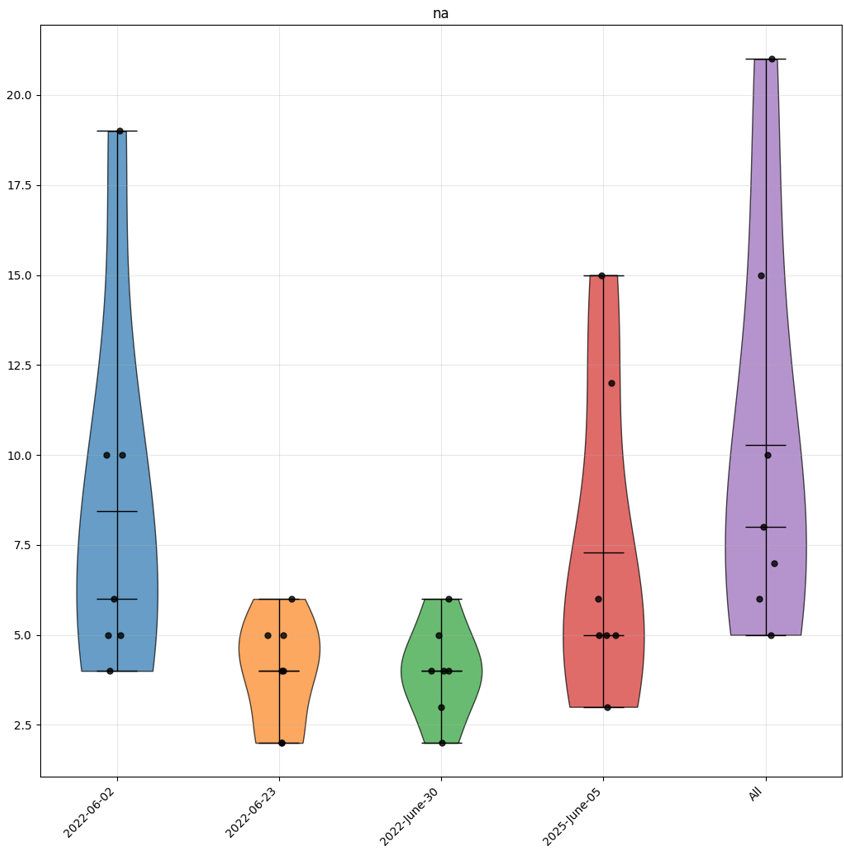

Figure S2a. Violin plots of the number of raw alleles (*NA*) across sampling dates.

**Fig. S2b**

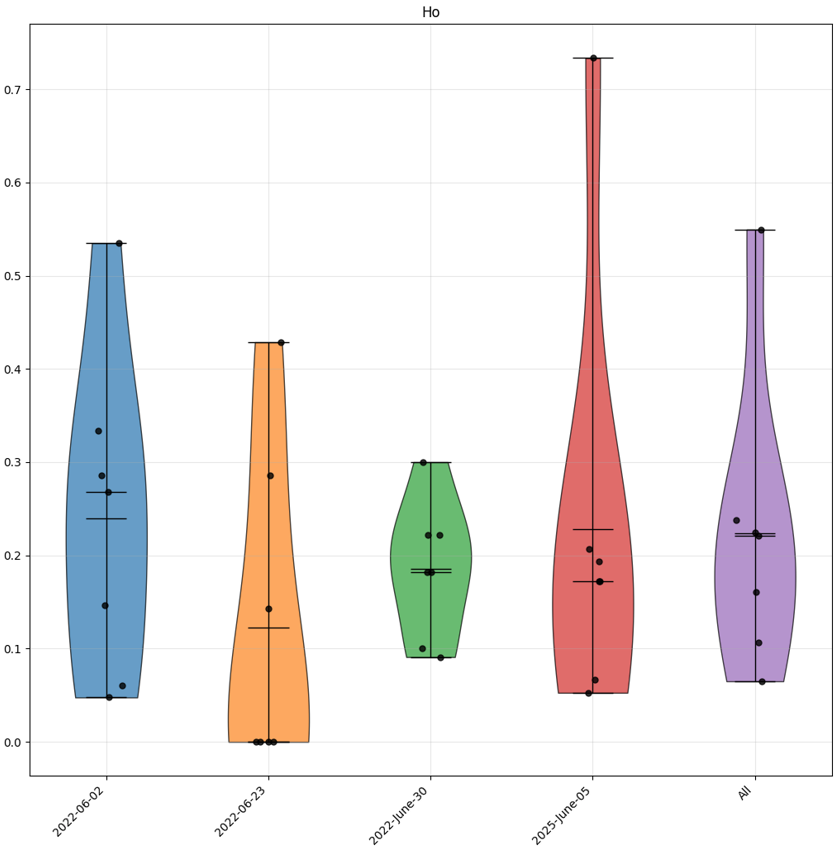

Figure S2b. Violin plots of the observed heterozygosity (*H_O_*) across sampling dates.

**Fig. S2c**

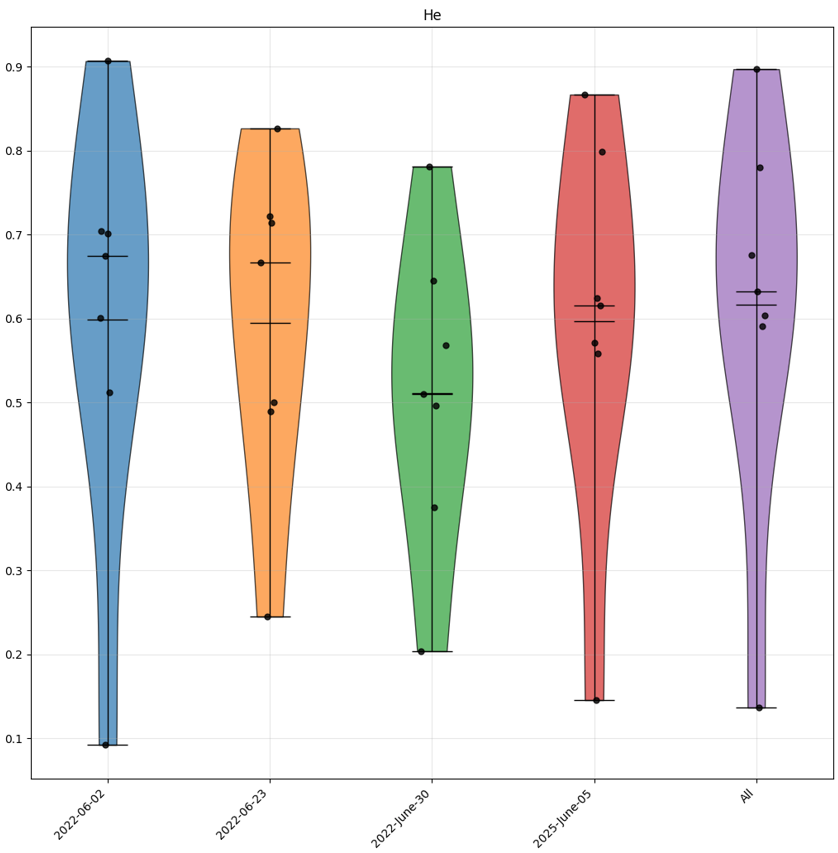

Figure S2c. Violin plots of the expected heterozygosity (*H_E_*) across sampling dates.

**Fig. S2d**

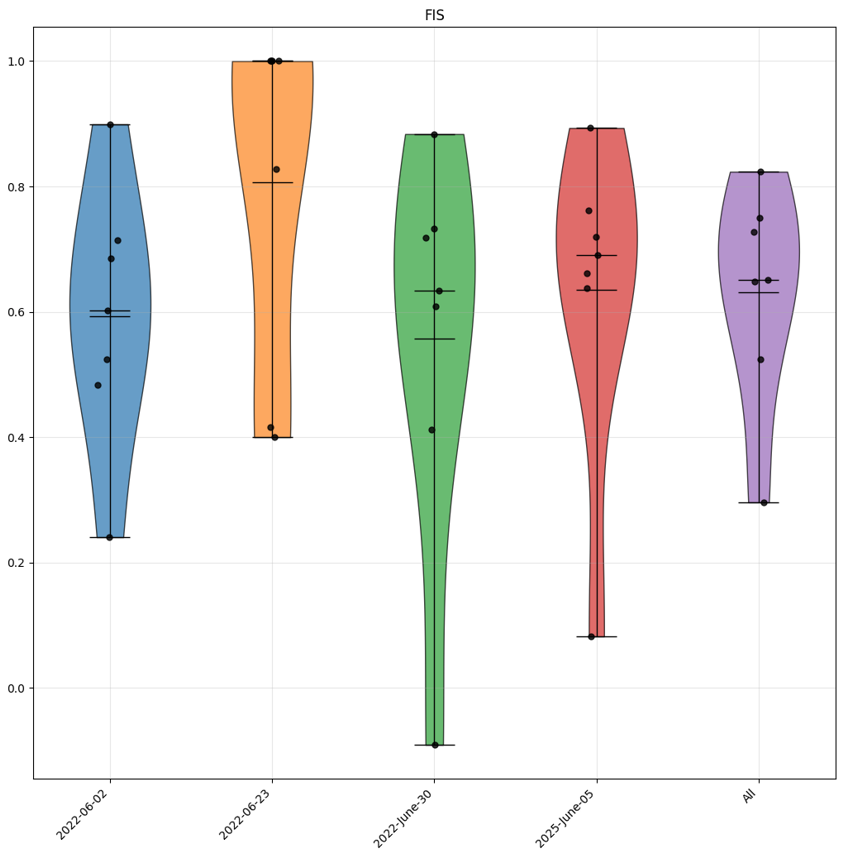

Figure S2d. Violin plots of the inbreeding coefficient (*F_IS_*) across sampling dates.

**Fig. S3**

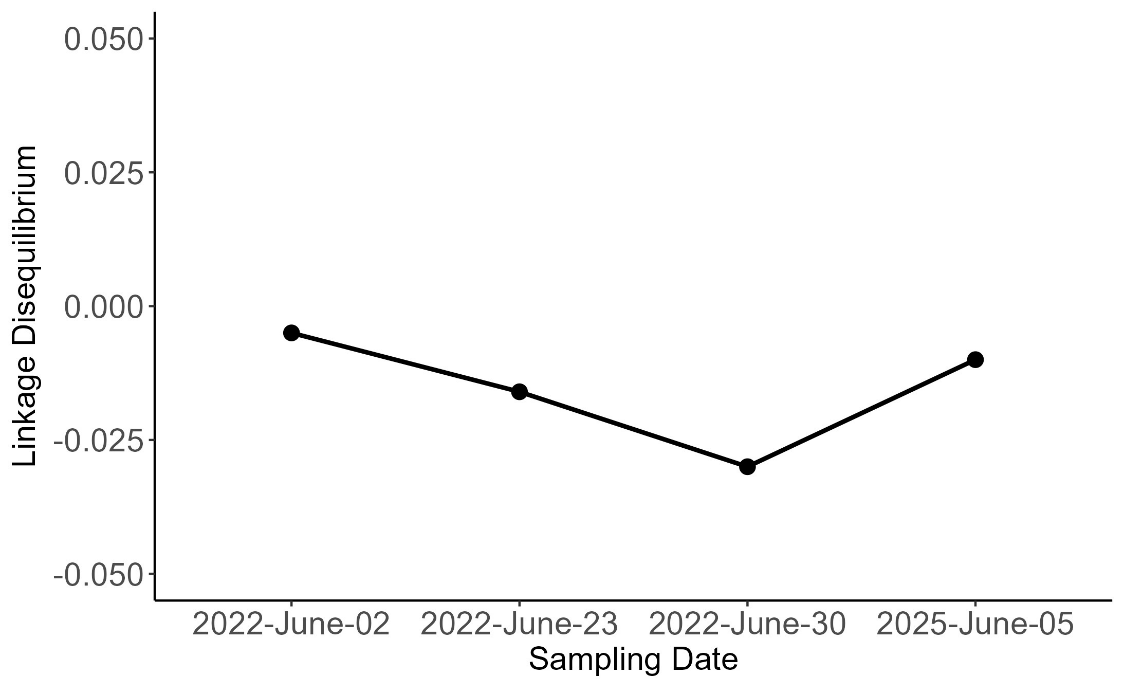

Figure S3. The trend in linkage disequilibrium (*r̄_D_*) values shown by each sampling date. Linkage disequilibrium ranges from -0.030 to -0.005 across sampling dates.

**Fig. S4**

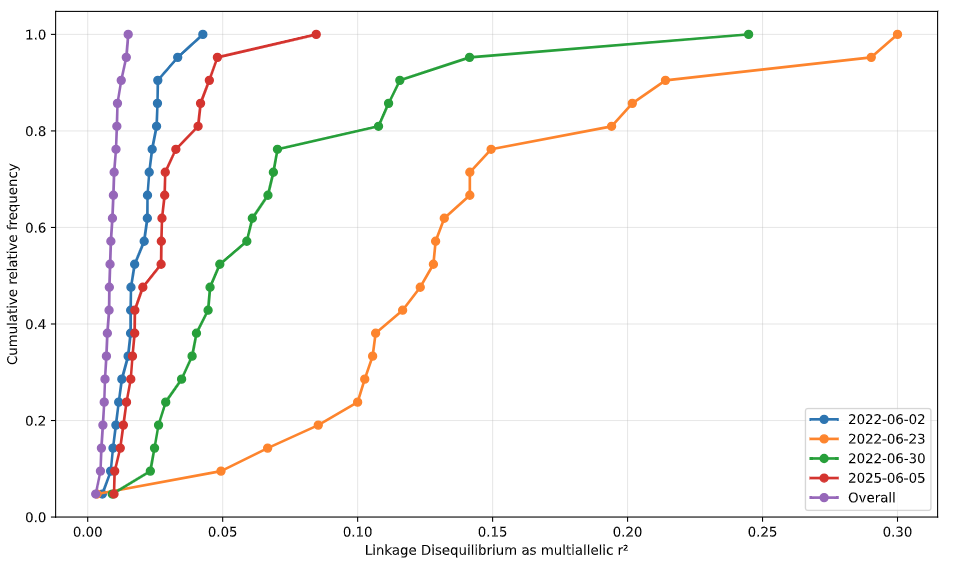

Figure S4. Plot of pairwise linkage disequilibrium by locus using Hill multiallelic r^2^ across each sampling date, as well as over each sampling date combined.

**Tables**

**Table S1** Metadata and sample size for sampled patches of *Chlainomonas* sp. All sampling areas were located on Upper Bagley Lake near Mount Baker in Washington state USA. Sampling area, letter tag used to identify area of snow field during sampling; latitude; longitude; sampling dates, dates on which each area was sampled; patches, names of individual patches of the *Chlainomonas* sp. bloom that were sampled at each area; cells extracted, the number of cells on which PicoPLEX whole genome amplification was conducted; cells partially amplified, the number of cells which had alleles on at a minimum of one locus; cells included in final dataset, the number of cells which had alleles on at least five loci and were used in downstream analyses.

| **Sampling area** | **Latitude** | **Longitude** | **Sampling dates** | **Patches** | **Cells extracted** | **Cells partially amplified** | **Cells included in final dataset** |
| --- | --- | --- | --- | --- | --- | --- | --- |
| A | 48.85443 | -121.69127 | 2 June 2022 | A1, A2, A7 | 5 | 5 | 3 |
|  |  |  | 23 June 2022 | A8, A9, A10 | 6 | 4 | 3 |
|  |  |  | 7 July 2022 | A8, A11 | 7 | 7 | 0 |
| B | 48.85440 | -121.69200 | 2 June 2022 | B1, B2, B3 | 5 | 5 | 4 |
|  |  |  | 30 June 2022 | B10, B11, B12 | 6 | 4 | 1 |
|  |  |  | 7 July 2022 | B12, B13, B14 | 6 | 3 | 1 |
| C | 48.85436 | -121.69230 | 5 June 2025 | C3 | 80 | 46 | 31 |
| G | 48.85389 | -121.69177 | 2 June 2022 | G1, G2, G3 | 5 | 4 | 1 |
|  |  |  | 23 June 2022 | G5, G6, G7 | 5 | 5 | 2 |
|  |  |  | 30 June 2022 | G9, G10 | 4 | 4 | 2 |
|  |  |  | 7 July 2022 | G8, G11 | 4 | 4 | 0 |
| V | 48.85377 | -121.69175 | 30 June 2022 | V2, V3 | 3 | 3 | 1 |
|  |  |  | 7 July 2022 | V1, V3, V4 | 6 | 6 | 2 |
| W | 48.85406 | -121.69127 | 23 June 2022 | W1, W2 | 3 | 2 | 1 |
|  |  |  | 30 June 2022 | W4 | 3 | 3 | 2 |
|  |  |  | 7 July 2022 | W2, W6, W7 | 5 | 3 | 0 |
| X | 48.85452 | -121.69153 | 2 June 2022 | X1, X2, X3 | 30 | 30 | 25 |
|  |  |  | 23 June 2022 | X3, X4, X5 | 3 | 3 | 1 |
|  |  |  | 30 June 2022 | X5, X6, X7 | 4 | 3 | 1 |
|  |  |  | 7 July 2022 | X6, X8, X9 | 5 | 1 | 0 |
| Y | 48.85452 | -121.69246 | 2 June 2022 | Y1, Y2, Y3 | 20 | 18 | 7 |
|  |  |  | 23 June 2022 | Y7 | 1 | 0 | 0 |
|  |  |  | 30 June 2022 | Y7, Y9, Y11 | 5 | 5 | 4 |
|  |  |  | 7 July 2022 | Y9, Y11, Y12 | 5 | 2 | 0 |
| Z | 48.85461 | -121.69260 | 2 June 2022 | Z2, Z3 | 9 | 7 | 4 |
|  |  |  | 23 June 2022 | Z5 | 1 | 1 | 0 |

**Table S2** Microsatellite genotypes of *Chlainomonas* sp. cells.

| **Patch** | **Date** | **Sample** | **Ch08** | | **Ch09** | | **Ch43** | | **Ch45** | | **Ch54** | | **Ch60** | | **Ch63** | |
| --- | --- | --- | --- | --- | --- | --- | --- | --- | --- | --- | --- | --- | --- | --- | --- | --- |
| X1 | 2022-June-02 | P3_X1_2 | 101 | 101 | 320 | 320 | 166 | 166 | 128 | 128 | 245 | 245 | 0 | 0 | 237 | 246 |
| X2 | 2022-June-02 | P3_X2_2 | 115 | 115 | 350 | 350 | 0 | 0 | 128 | 128 | 237 | 241 | 148 | 148 | 0 | 0 |
| X2 | 2022-June-02 | P3_X2_3 | 101 | 101 | 304 | 304 | 166 | 166 | 128 | 128 | 241 | 281 | 148 | 148 | 237 | 246 |
| X1 | 2022-June-02 | P3_X1_5 | 97 | 101 | 304 | 304 | 166 | 166 | 0 | 0 | 241 | 241 | 0 | 0 | 246 | 246 |
| X2 | 2022-June-02 | P3_X2_5 | 101 | 101 | 306 | 306 | 166 | 166 | 128 | 128 | 241 | 251 | 124 | 124 | 246 | 246 |
| X1 | 2022-June-02 | P3_X1_6 | 101 | 101 | 310 | 310 | 166 | 166 | 128 | 128 | 237 | 237 | 148 | 148 | 0 | 0 |
| X2 | 2022-June-02 | P3_X2_6 | 97 | 97 | 316 | 316 | 166 | 166 | 128 | 128 | 241 | 281 | 151 | 151 | 237 | 237 |
| X1 | 2022-June-02 | P3_X1_7 | 97 | 101 | 320 | 320 | 166 | 166 | 128 | 128 | 237 | 277 | 148 | 148 | 243 | 243 |
| X2 | 2022-June-02 | P3_X2_7 | 101 | 101 | 316 | 316 | 166 | 166 | 120 | 120 | 241 | 281 | 148 | 148 | 246 | 246 |
| X1 | 2022-June-02 | P3_X1_8 | 101 | 115 | 300 | 300 | 166 | 166 | 128 | 128 | 237 | 251 | 148 | 148 | 237 | 246 |
| X2 | 2022-June-02 | P3_X2_8 | 115 | 115 | 308 | 320 | 166 | 166 | 128 | 128 | 241 | 241 | 148 | 148 | 237 | 237 |
| X3 | 2022-June-02 | P3_X3_8 | 97 | 97 | 314 | 326 | 166 | 166 | 120 | 128 | 237 | 241 | 148 | 148 | 243 | 249 |
| Y1 | 2022-June-02 | P3_Y1_8 | 97 | 101 | 308 | 308 | 166 | 166 | 128 | 128 | 0 | 0 | 124 | 124 | 246 | 246 |
| X1 | 2022-June-02 | P3_X1_9 | 101 | 101 | 304 | 314 | 166 | 166 | 128 | 128 | 237 | 277 | 145 | 145 | 246 | 246 |
| X2 | 2022-June-02 | P3_X2_10 | 101 | 101 | 318 | 318 | 166 | 166 | 120 | 120 | 241 | 241 | 148 | 148 | 0 | 0 |
| X1 | 2022-June-02 | P3_X1_11 | 101 | 101 | 314 | 314 | 166 | 166 | 120 | 128 | 241 | 281 | 148 | 148 | 246 | 246 |
| X2 | 2022-June-02 | P3_X2_11 | 97 | 101 | 0 | 0 | 166 | 166 | 124 | 128 | 241 | 281 | 148 | 148 | 246 | 246 |
| Y2 | 2022-June-02 | P3_Y2B_11 | 97 | 101 | 320 | 320 | 166 | 166 | 124 | 124 | 241 | 241 | 124 | 124 | 246 | 246 |
| X1 | 2022-June-02 | P3_X1_12 | 101 | 115 | 308 | 308 | 166 | 166 | 128 | 128 | 237 | 277 | 148 | 148 | 246 | 246 |
| X2 | 2022-June-02 | P3_X2_12 | 97 | 101 | 310 | 310 | 166 | 166 | 128 | 128 | 239 | 239 | 124 | 124 | 249 | 249 |
| C3 | 2025-June-05 | C3_84 | 101 | 101 | 312 | 312 | 166 | 166 | 128 | 128 | 241 | 281 | 124 | 148 | 246 | 246 |
| C3 | 2025-June-05 | C3_85 | 111 | 111 | 312 | 312 | 166 | 166 | 128 | 128 | 241 | 241 | 124 | 145 | 246 | 246 |
| A7 | 2022-June-02 | P4_A7_3 | 101 | 101 | 312 | 320 | 166 | 166 | 0 | 0 | 237 | 237 | 124 | 124 | 246 | 246 |
| B3 | 2022-June-02 | P4_B3_3 | 101 | 101 | 316 | 316 | 166 | 166 | 128 | 128 | 241 | 241 | 0 | 0 | 243 | 243 |
| B1 | 2022-June-02 | P4_B1_4 | 111 | 111 | 312 | 312 | 166 | 166 | 128 | 128 | 241 | 245 | 124 | 148 | 243 | 243 |
| B2 | 2022-June-02 | P4_B2_4 | 0 | 0 | 336 | 336 | 166 | 166 | 120 | 120 | 241 | 241 | 124 | 124 | 240 | 243 |
| G1 | 2022-June-02 | P4_G1_4 | 0 | 0 | 312 | 324 | 166 | 166 | 124 | 124 | 237 | 277 | 124 | 148 | 243 | 246 |
| A1 | 2022-June-02 | P4_A1_5 | 97 | 101 | 304 | 316 | 166 | 166 | 120 | 120 | 237 | 241 | 124 | 124 | 243 | 243 |
| A2 | 2022-June-02 | P4_A2_5 | 0 | 0 | 324 | 324 | 166 | 166 | 128 | 128 | 237 | 241 | 0 | 0 | 243 | 243 |
| B3 | 2022-June-02 | P4_B3_5 | 0 | 0 | 290 | 310 | 166 | 166 | 128 | 128 | 241 | 241 | 124 | 124 | 246 | 246 |
| X3 | 2022-June-02 | P4_X3_1 | 0 | 0 | 0 | 0 | 166 | 166 | 120 | 120 | 237 | 237 | 124 | 124 | 243 | 243 |
| Y2 | 2022-June-02 | P4_Y2_1 | 0 | 0 | 304 | 312 | 166 | 166 | 120 | 128 | 237 | 277 | 124 | 124 | 246 | 246 |
| X1 | 2022-June-02 | P4_X1_2 | 103 | 103 | 362 | 362 | 166 | 166 | 120 | 128 | 241 | 241 | 0 | 0 | 243 | 243 |
| X1 | 2022-June-02 | P4_X1_3 | 97 | 97 | 312 | 312 | 166 | 174 | 120 | 120 | 237 | 237 | 124 | 124 | 249 | 249 |
| X1 | 2022-June-02 | P4_X1_4 | 117 | 117 | 306 | 306 | 166 | 166 | 0 | 0 | 241 | 251 | 0 | 0 | 246 | 246 |
| X3 | 2022-June-02 | P4_X3_4 | 0 | 0 | 288 | 306 | 166 | 166 | 120 | 120 | 243 | 243 | 124 | 124 | 246 | 249 |
| Y1 | 2022-June-02 | P4_Y1_4 | 0 | 0 | 304 | 304 | 166 | 166 | 128 | 128 | 231 | 237 | 124 | 124 | 243 | 243 |
| Y3 | 2022-June-02 | P4_Y3_4 | 103 | 103 | 306 | 326 | 0 | 0 | 128 | 128 | 241 | 251 | 0 | 0 | 246 | 246 |
| X2 | 2022-June-02 | P4_X2_5 | 105 | 109 | 310 | 310 | 166 | 166 | 128 | 128 | 241 | 281 | 0 | 0 | 237 | 237 |
| X3 | 2022-June-02 | P4_X3_5 | 123 | 123 | 310 | 310 | 166 | 166 | 128 | 128 | 237 | 237 | 124 | 124 | 237 | 237 |
| Y1 | 2022-June-02 | P4_Y1_5 | 0 | 0 | 312 | 312 | 166 | 166 | 124 | 124 | 241 | 251 | 145 | 145 | 243 | 246 |
| Y2 | 2022-June-02 | P4_Y2_5 | 0 | 0 | 296 | 310 | 166 | 166 | 120 | 128 | 241 | 241 | 145 | 145 | 243 | 246 |
| Z2 | 2022-June-02 | P4_Z2_2 | 0 | 0 | 312 | 312 | 164 | 164 | 128 | 128 | 241 | 241 | 0 | 0 | 243 | 249 |
| Z2 | 2022-June-02 | P4_Z2_4 | 0 | 0 | 312 | 312 | 166 | 166 | 120 | 120 | 241 | 241 | 124 | 124 | 237 | 243 |
| Z2 | 2022-June-02 | P4_Z2_6 | 0 | 0 | 312 | 312 | 160 | 166 | 128 | 128 | 237 | 241 | 0 | 0 | 246 | 246 |
| Z3 | 2022-June-02 | P4_Z3_6 | 0 | 0 | 308 | 316 | 166 | 166 | 120 | 120 | 241 | 241 | 0 | 0 | 243 | 243 |
| A8 | 2022-June-23 | T4_A8_5 | 97 | 101 | 308 | 308 | 166 | 166 | 116 | 116 | 241 | 281 | 0 | 0 | 246 | 246 |
| A9 | 2022-June-23 | T4_A9_1 | 101 | 101 | 314 | 314 | 166 | 166 | 124 | 124 | 277 | 277 | 0 | 0 | 0 | 0 |
| A9 | 2022-June-23 | T4_A9_2 | 97 | 101 | 314 | 324 | 164 | 164 | 112 | 112 | 277 | 277 | 145 | 145 | 246 | 246 |
| X4 | 2022-June-23 | T4_X4_3 | 97 | 97 | 300 | 300 | 166 | 166 | 128 | 128 | 281 | 281 | 124 | 124 | 246 | 246 |
| W2 | 2022-June-23 | T4_W2_1 | 97 | 97 | 324 | 324 | 166 | 166 | 128 | 128 | 269 | 277 | 148 | 148 | 243 | 243 |
| G5 | 2022-June-23 | T4_G5_4 | 101 | 101 | 318 | 318 | 166 | 166 | 0 | 0 | 241 | 241 | 0 | 0 | 246 | 246 |
| G5 | 2022-June-23 | T4_G5_5 | 101 | 101 | 320 | 320 | 166 | 166 | 116 | 116 | 237 | 277 | 0 | 0 | 237 | 237 |
| X7 | 2022-June-30 | T5_X7_3 | 97 | 101 | 312 | 312 | 166 | 166 | 120 | 128 | 281 | 281 | 148 | 148 | 0 | 0 |
| G9 | 2022-June-30 | T5_G9_1 | 101 | 101 | 320 | 320 | 166 | 166 | 0 | 0 | 281 | 281 | 148 | 148 | 243 | 243 |
| V2 | 2022-June-30 | T5_V2_1 | 101 | 101 | 304 | 304 | 0 | 0 | 128 | 128 | 281 | 281 | 148 | 148 | 243 | 243 |
| Y9 | 2022-June-30 | T5_Y9_2 | 97 | 97 | 312 | 312 | 166 | 166 | 120 | 120 | 241 | 241 | 124 | 124 | 243 | 243 |
| Y9 | 2022-June-30 | T5_Y9_3 | 97 | 97 | 312 | 312 | 166 | 166 | 120 | 120 | 277 | 277 | 148 | 148 | 243 | 246 |
| G10 | 2022-June-30 | T5_G10_3 | 97 | 97 | 310 | 310 | 166 | 166 | 0 | 0 | 241 | 281 | 0 | 0 | 243 | 243 |
| Y11 | 2022-June-30 | T5_Y11_1 | 97 | 97 | 304 | 304 | 166 | 166 | 128 | 128 | 277 | 277 | 148 | 148 | 240 | 243 |
| Y11 | 2022-June-30 | T5_Y11_3 | 97 | 97 | 308 | 324 | 166 | 166 | 120 | 120 | 281 | 281 | 124 | 148 | 246 | 246 |
| W4 | 2022-June-30 | T5_W4_2 | 101 | 101 | 320 | 320 | 0 | 0 | 120 | 120 | 277 | 277 | 124 | 124 | 246 | 246 |
| W4 | 2022-June-30 | T5_W4_3 | 101 | 101 | 304 | 304 | 162 | 166 | 116 | 116 | 237 | 277 | 148 | 148 | 243 | 243 |
| B11 | 2022-June-30 | T5_B11_1 | 97 | 101 | 310 | 310 | 152 | 166 | 120 | 128 | 277 | 277 | 148 | 148 | 237 | 243 |
| V4 | 2022-July-07 | T6_V4_4 | 97 | 97 | 360 | 360 | 0 | 0 | 128 | 128 | 279 | 279 | 148 | 148 | 243 | 243 |
| B12 | 2022-July-07 | T6_B12_3 | 97 | 97 | 0 | 0 | 166 | 166 | 0 | 0 | 281 | 281 | 148 | 148 | 246 | 246 |
| V1 | 2022-July-07 | T6_V1_1 | 97 | 97 | 310 | 322 | 166 | 166 | 128 | 128 | 277 | 281 | 148 | 148 | 243 | 243 |
| C3 | 2025-June-05 | C3_1 | 0 | 0 | 312 | 312 | 166 | 166 | 128 | 128 | 251 | 293 | 124 | 124 | 237 | 243 |
| C3 | 2025-June-05 | C3_4 | 97 | 97 | 310 | 316 | 166 | 166 | 128 | 128 | 243 | 285 | 124 | 124 | 243 | 243 |
| C3 | 2025-June-05 | C3_5 | 97 | 101 | 300 | 300 | 166 | 166 | 116 | 128 | 241 | 281 | 148 | 148 | 246 | 246 |
| C3 | 2025-June-05 | C3_6 | 0 | 0 | 306 | 306 | 166 | 166 | 124 | 128 | 243 | 285 | 148 | 148 | 237 | 237 |
| C3 | 2025-June-05 | C3_7 | 0 | 0 | 304 | 304 | 166 | 166 | 128 | 128 | 239 | 279 | 124 | 148 | 243 | 246 |
| C3 | 2025-June-05 | C3_9 | 0 | 0 | 0 | 0 | 166 | 166 | 124 | 124 | 241 | 281 | 145 | 145 | 246 | 246 |
| C3 | 2025-June-05 | C3_11 | 0 | 0 | 310 | 310 | 166 | 166 | 116 | 116 | 237 | 277 | 148 | 148 | 0 | 0 |
| C3 | 2025-June-05 | C3_13 | 0 | 0 | 296 | 312 | 166 | 166 | 128 | 128 | 241 | 241 | 124 | 145 | 246 | 246 |
| C3 | 2025-June-05 | C3_15 | 97 | 97 | 308 | 324 | 172 | 172 | 120 | 120 | 257 | 257 | 148 | 148 | 246 | 246 |
| C3 | 2025-June-05 | C3_16 | 101 | 101 | 316 | 316 | 166 | 166 | 128 | 128 | 241 | 281 | 145 | 145 | 243 | 243 |
| C3 | 2025-June-05 | C3_19 | 0 | 0 | 314 | 314 | 166 | 166 | 120 | 120 | 241 | 281 | 148 | 148 | 246 | 246 |
| C3 | 2025-June-05 | C3_27 | 0 | 0 | 316 | 316 | 0 | 0 | 112 | 112 | 237 | 277 | 124 | 124 | 246 | 246 |
| C3 | 2025-June-05 | C3_28 | 0 | 0 | 350 | 350 | 0 | 0 | 124 | 128 | 241 | 281 | 148 | 148 | 243 | 246 |
| C3 | 2025-June-05 | C3_31 | 97 | 97 | 312 | 312 | 0 | 0 | 128 | 128 | 239 | 239 | 148 | 148 | 246 | 246 |
| C3 | 2025-June-05 | C3_35 | 0 | 0 | 314 | 314 | 0 | 0 | 120 | 120 | 241 | 281 | 148 | 148 | 246 | 246 |
| C3 | 2025-June-05 | C3_38 | 0 | 0 | 312 | 312 | 0 | 0 | 120 | 120 | 237 | 277 | 124 | 124 | 243 | 243 |
| C3 | 2025-June-05 | C3_39 | 97 | 97 | 302 | 312 | 0 | 0 | 120 | 128 | 241 | 281 | 124 | 124 | 243 | 246 |
| C3 | 2025-June-05 | C3_41 | 97 | 97 | 316 | 316 | 0 | 0 | 128 | 128 | 241 | 241 | 0 | 0 | 246 | 246 |
| C3 | 2025-June-05 | C3_48 | 97 | 97 | 310 | 310 | 166 | 166 | 128 | 128 | 237 | 277 | 151 | 151 | 243 | 243 |
| C3 | 2025-June-05 | C3_51 | 0 | 0 | 302 | 320 | 166 | 166 | 128 | 128 | 241 | 241 | 124 | 124 | 243 | 243 |
| C3 | 2025-June-05 | C3_53 | 97 | 97 | 308 | 320 | 0 | 0 | 124 | 124 | 237 | 277 | 148 | 148 | 243 | 243 |
| C3 | 2025-June-05 | C3_56 | 0 | 0 | 330 | 330 | 0 | 0 | 120 | 120 | 237 | 277 | 148 | 148 | 246 | 246 |
| C3 | 2025-June-05 | C3_57 | 0 | 0 | 320 | 320 | 166 | 172 | 128 | 128 | 237 | 277 | 0 | 0 | 246 | 246 |
| C3 | 2025-June-05 | C3_60 | 105 | 105 | 312 | 312 | 0 | 0 | 124 | 128 | 241 | 241 | 148 | 148 | 0 | 0 |
| C3 | 2025-June-05 | C3_65 | 97 | 97 | 312 | 312 | 0 | 0 | 128 | 128 | 241 | 281 | 148 | 148 | 243 | 243 |
| C3 | 2025-June-05 | C3_66 | 103 | 103 | 304 | 304 | 166 | 166 | 128 | 128 | 241 | 281 | 124 | 124 | 243 | 243 |
| C3 | 2025-June-05 | C3_75 | 0 | 0 | 0 | 0 | 166 | 166 | 128 | 128 | 241 | 241 | 124 | 124 | 243 | 243 |
| C3 | 2025-June-05 | C3_79 | 103 | 103 | 310 | 310 | 166 | 166 | 128 | 128 | 0 | 0 | 148 | 148 | 246 | 253 |
| C3 | 2025-June-05 | C3_83 | 0 | 0 | 320 | 320 | 0 | 0 | 120 | 128 | 237 | 251 | 145 | 148 | 243 | 243 |

**Table S3** Primer development for Chlainomonas sp., showing results for each locus with both unlabeled oligos and on the sequencer.

| **Locus** | **Forward Sequence** | **Reverse Sequence** | **Motif** | **Expected Size** | **Unlabeled Oligo Tests** | **Dye** | **Sequencer Results** |
| --- | --- | --- | --- | --- | --- | --- | --- |
| Ch01 | GAGACCGGGAGCGAAGATAC | TCTGATGCCTCGTCTTGGAG | Di | 195 | Single band | 6FAM | No peaks |
| Ch02 | TCGTCTTCTCCTTCACTGCC | GAGAGGGAGAGGTGTCGATG | Di | 251 | Single band | VIC | Too noisy to score |
| Ch03 | CGCTCAGCTGTAATCACGAC | CTGAAGACGCATTGAAGGGC | Tri | 373 | Single band | PET | No peaks |
| Ch04 | GACAACCGTTATGTGCAGCC | CCACTCACCATGATGAAGCG | Tri | 306 | Single band | NED | Inconsistent amplification |
| Ch05 | GCGCCATCAATTTCCTCCTC | CGCGGTAGACTTGGTTTCAC | Di | 323 | No amplification | Not tested on capillary sequencer | - |
| Ch06 | AAACATCAGCCGCGGTTAAG | GTCTCCAGCACCGAGTAGAG | Di | 328 | Multiband profile | Not tested on capillary sequencer | - |
| Ch07 | CCAAGGAGAGTCTGAGGAGC | AAGATGCGGATTTGTGGCTG | Di | 176 | No amplification | Not tested on capillary sequencer | - |
| Ch08 | AAAGGGTGTTGGAACTGTGC | TTTGCTGGCTACTCGAGGTC | Di | 176 | Single band | 6FAM | Used in analysis |
| Ch09 | TGGTTTGCGGTCATTGTGTC | CGTAGGTGTCATGGTGCAAC | Di | 307 | Single band | NED | Used in analysis |
| Ch10 | AACCACATCAAGCAAGCGAG | TCGTGCTTACAAACACTGGC | Di | 184 | Single band | 6FAM | Too noisy to score |
| Ch11 | CAGCGAAGACTTGAAGCCAG | TCGCCGATTTAGACCTCCTG | Tri | 218 | Single band | VIC | Too noisy to score |
| Ch12 | ACAATGCAGCAACAGTCAGG | GCTGCCGTCGTCATCTTTAC | Di | 238 | Single band | VIC | Too noisy to score |
| Ch13 | GTTGCTCCGTATTGTGGCTG | CTCGCGTTCATGTTTCCTCC | Di | 436 | Single band | PET | No peaks |
| Ch14 | GCAGAATCACCACAGCCATC | ATTCGCACTTAGTCGCTTGC | Di | 401 | Single band | PET | Too noisy to score |
| Ch15 | GGTAGGCTGGTATCGTGAGG | GAGAACGTGCCATGTGAGAC | Di | 212 | Multiband profile | Not tested on capillary sequencer | - |
| Ch16 | GCGGAGAAGTTCGTTGTGAG | GCAAGCAGAGAGTCAAGAGC | Tetra | 175 | Multiband profile | Not tested on capillary sequencer | - |
| Ch17 | GACAGGTAGCCAGAGTCCAC | TGTGGCTGGAAGTAGGACTG | Di | 236 | Multiband profile | Not tested on capillary sequencer | - |
| Ch18 | TCCTCCGAATGTCACCACTC | GAAGACCGTCATTTGGCCTG | Tetra | 263 | Single band | NED | Too noisy to score |
| Ch19 | CGCGAGACCAAGAACTGATG | GAAGTCGAGGTGGGAGAGTC | Di | 368 | Single band | PET | Inconsistent amplification |
| Ch20 | AATGCATCAGTAATCGCCGG | CTAGGCGGTGCACATTTCAG | Di | 117 | No amplification | Not tested on capillary sequencer | - |
| Ch21 | GCTGCACACTCTGTTACCTG | CTCTACCATCCGCCCTTCTC | Di | 181 | Single band | 6FAM | No peaks |
| Ch22 | TGTGCAGGTAGGAGCTTGAG | CGTGACGTCCTTAAGCAACG | Tetra | 378 | Single band | NED | Too noisy to score |
| Ch23 | AGCAGTCCACAGGTGTAGTG | TCAAGAAACAAGACGCTCGC | Di | 272 | Single band | VIC | Too noisy to score |
| Ch24 | ATTGACGAGGTTGTGCTTGC | AGACACGAAACCATCCTCCC | Di | 303 | Single band | NED | Too noisy to score |
| Ch25 | GCATCAACTCACGACCACAG | CTGTATTTGGAGCCGCTGAC | Tetra | 302 | Multiband profile | Not tested on capillary sequencer | - |
| Ch26 | GCTGGTATCCACAACCTTGC | TGCTGGACACTGGCCTATAC | Tetra | 324 | Single band | NED | Too noisy to score |
| Ch27 | GTACAGCTGGACATGGTTGC | TCCTCATTGCCTGCTCTCTG | Di | 123 | Multiband profile | Not tested on capillary sequencer | - |
| Ch28 | CCTTACGCATGCACACTCAG | GCTGCACACGAACGATACAG | Di | 160 | Single band | 6FAM | Monomorphic |
| Ch29 | TGCTCCCTCTTGTGACTGTC | TGCCGTTTGATCAGCATGTG | Tetra | 403 | No amplification | Not tested on capillary sequencer | - |
| Ch30 | CTCAGCGCTTCATTCTCTGC | AATCCCGTAGAAGTAGCCGG | Di | 206 | Single band | VIC | Too noisy to score |
| Ch31 | CTTGTGTCCTGTGTCCTTGC | CAGCACACAGACATCGGATG | Di | 410 | Single band | PET | No peaks |
| Ch32 | TTGTCCTGTTGCGCTGATTG | CGTCTTCGTTGTTGGGATGG | Di | 253 | Multiband profile | Not tested on capillary sequencer | - |
| Ch33 | TCTCCTTCCCGCACATACAG | CGACTTCCCAACGATCAAGC | Di | 353 | Single band | PET | No peaks |
| Ch34 | TCGCCTTGAGTCCTATGTCG | TAACGACCACCCATCACGAG | Di | 228 | Single band | VIC | Too noisy to score |
| Ch35 | GCCTCTGTGCAGGTATTTCG | AGTCATCGCAGCCAATTTCG | Tetra | 361 | Single band | NED | Too noisy to score |
| Ch36 | AATGGCATGCGATCAAGTGG | CGCAAGTCGTCTGGTTTCTC | Di | 389 | Multiband profile | Not tested on capillary sequencer | - |
| Ch37 | AATGCACGTGTCGGAGTTTG | CGAGTCTGTGAGAGGTGGAG | Tetra | 174 | Multiband profile | Not tested on capillary sequencer | - |
| Ch38 | TGTGACTCCATGCTCTACGG | ACAACCAGGAACAATGCAGC | Di | 287 | Single band | NED | Too noisy to score |
| Ch39 | TGCTTGCCAGTCTTGCTTTC | GAGGGTCGCGAATTCAAACC | Di | 162 | Single band | 6FAM | No peaks |
| Ch40 | TTCCTGCCCTCTCCATGTAC | GTCAGATGCAGGCGAGAAAC | Di | 248 | Multiband profile | Not tested on capillary sequencer | - |
| Ch41 | AAACAAACGACCAACCGACC | AAGCACGCAACACTTAGCAG | Di | 292 | Multiband profile | Not tested on capillary sequencer | - |
| Ch42 | ACAGCTCCAGAGAAGCAGTC | GCAGCTCACACACTGATGAC | Tri | 242 | No amplification | Not tested on capillary sequencer | - |
| Ch43 | ACAACCCGAGTATGTCCCTC | AATACACCTCGTCGTCCCAG | Di | 187 | Single band | 6FAM | Used in analysis |
| Ch44 | TGTGGCATAGGAGGAGCATC | TACCCTGCCAAATCCCAGTC | Di | 338 | No amplification | Not tested on capillary sequencer | - |
| Ch45 | CCACAAACCAACTTCCCACC | TGACGCTCCTGGAGAATGAG | Tetra | 131 | Single band | 6FAM | Used in analysis |
| Ch46 | TATACACTCACCCGTCGCTG | AGCATGCTTTCGGACCTTTG | Tetra | 193 | No amplification | Not tested on capillary sequencer | - |
| Ch47 | GGTTGAGCGGATTCAGGTTG | AGCACACCTCTCCAGTTCAG | Di | 109 | No amplification | Not tested on capillary sequencer | - |
| Ch48 | TGGTGGCTCGTGATCTCTAC | GATCGAAACCTGACCAAGCG | Di | 233 | Single band | VIC | No peaks |
| Ch49 | TAGACATCGGGTGCGAGATG | ATTCCTGCGTAGTCCCAGTG | Di | 295 | No amplification | Not tested on capillary sequencer | - |
| Ch50 | GAAAGTGGACAGCAAGCGAG | CTGCACCCGAATGTCTGTTG | Tetra | 277 | No amplification | Not tested on capillary sequencer | - |
| Ch51 | TGTATTCCCAGGCAGCTCAG | ACCGTGCACTACCTGATACC | Di | 227 | No amplification | Not tested on capillary sequencer | - |
| Ch52 | CTGAAACTGGTAGGCAAGCG | GCAAGTCAGAGAAAGCGGAC | Tri | 108 | Multiband profile | Not tested on capillary sequencer | - |
| Ch53 | GACAGGGCAATAGAAACGGC | GGCATGCTTCTACGGTTGAC | Di | 240 | Single band | NED | Too noisy to score |
| Ch54 | GCGGGTCAGTTTGTGTGTAC | AAAGGCGAGTCGATTTGCAG | Di | 237 | Single band | PET | Used in analysis |
| Ch55 | AATGATGCAACGCCTCGAAG | AGGTGATGCTACGGTGCTAG | Di | 382 | Single band | 6FAM | Inconsistent amplification |
| Ch56 | AGCCGATACACTAGTGCAGG | TTTCCTCCACTCCACCAGAC | Di | 282 | No amplification | Not tested on capillary sequencer | - |
| Ch57 | TGGGAAAGCTTGGGTCTCTC | TCAATGGAGTGGATGGAGGC | Di | 241 | Single band | VIC | Too noisy to score |
| Ch58 | AACACAAATCAGGCAGCCAC | GGTGGACAAGCGGGAAATAC | Di | 183 | Single band | NED | Too noisy to score |
| Ch59 | TGTCTGTATCTGTCGGGCTG | TGACCATACACGTACCACCG | Di | 112 | Multiband profile | Not tested on capillary sequencer | - |
| Ch60 | AGTTGGGTCAGACTCGTGAC | GCCCTTGTCTATTTGCACCC | Tri | 146 | Single band | PET | Used in analysis |
| Ch61 | GGCTACCACAACAGAAGCAC | ATGCATATCGAGAGGCCCTG | Di | 120 | Multiband profile | Not tested on capillary sequencer | - |
| Ch62 | GCTTCTTGCGAGCCTTCTTC | CAACGCAGGGATGTACACAC | Di | 347 | No amplification | Not tested on capillary sequencer | - |
| Ch63 | CAAGCAAGCAGACGTCTTCC | CGGTAGCTTCTGGTGTTTGC | Tri | 248 | Single band | 6FAM | Used in analysis |
| Ch64 | AATCCGTTTGCAGCAGTGAC | CCAGACATTCCAGCGTGTTC | Di | 138 | Single band | VIC | Inconsistent amplification |
| Ch65 | AAACGAGAGCACCAGAAAGC | GATGTTGTCGCGTGTCTCAG | Di | 219 | No amplification | Not tested on capillary sequencer | - |
| Ch66 | AGTTCCTCTTGCACTCGGAG | TGTGTCGGACCTCATACGTG | Di | 181 | No amplification | Not tested on capillary sequencer | - |
| Ch67 | GGAAGGACTGGAGCCATAGG | TAGGGTGTTGGTGATGGCTC | Di | 340 | No amplification | Not tested on capillary sequencer | - |

**Table S4** Allele call table showing the locus, dye, raw allele range, and final allele call. The raw calls are based on fragment analysis preformed on the SeqStudio24 at the Virginia Institute of Marine Science Eastern Shore Laboratory.

| **Locus** | **Motif** | **Dye** | **Raw allele score min** | **Raw allele score max** | **Call** |
| --- | --- | --- | --- | --- | --- |
| Ch08 | Di | 6-FAM | 90.833 | 90.833 | 91 |
| Ch08 | Di | 6-FAM | 94.909 | 94.909 | 95 |
| Ch08 | Di | 6-FAM | 97.523 | 98.195 | 97 |
| Ch08 | Di | 6-FAM | 98.778 | 99.385 | 99 |
| Ch08 | Di | 6-FAM | 101.302 | 102.251 | 101 |
| Ch08 | Di | 6-FAM | 102.959 | 103.233 | 103 |
| Ch08 | Di | 6-FAM | 105.771 | 106.14 | 105 |
| Ch08 | Di | 6-FAM | 107.857 | 107.857 | 107 |
| Ch08 | Di | 6-FAM | 109.082 | 109.263 | 109 |
| Ch08 | Di | 6-FAM | 112.398 | 112.419 | 111 |
| Ch08 | Di | 6-FAM | 113.374 | 113.374 | 113 |
| Ch08 | Di | 6-FAM | 114.924 | 115.024 | 115 |
| Ch08 | Di | 6-FAM | 116.153 | 116.629 | 117 |
| Ch08 | Di | 6-FAM | 123.204 | 123.204 | 123 |
| Ch09 | Di | NED | 286.565 | 286.565 | 286 |
| Ch09 | Di | NED | 288.35 | 289.176 | 288 |
| Ch09 | Di | NED | 290.057 | 290.057 | 290 |
| Ch09 | Di | NED | 294.109 | 294.109 | 294 |
| Ch09 | Di | NED | 296.317 | 296.742 | 296 |
| Ch09 | Di | NED | 299.336 | 299.336 | 298 |
| Ch09 | Di | NED | 300.282 | 301.148 | 300 |
| Ch09 | Di | NED | 301.502 | 302.072 | 302 |
| Ch09 | Di | NED | 304.025 | 304.914 | 304 |
| Ch09 | Di | NED | 305.521 | 306.627 | 306 |
| Ch09 | Di | NED | 308.103 | 309.116 | 308 |
| Ch09 | Di | NED | 310.005 | 310.771 | 310 |
| Ch09 | Di | NED | 311.81 | 312.687 | 312 |
| Ch09 | Di | NED | 313.237 | 314.892 | 314 |
| Ch09 | Di | NED | 315.833 | 316.802 | 316 |
| Ch09 | Di | NED | 317.772 | 318.562 | 318 |
| Ch09 | Di | NED | 320.105 | 321.111 | 320 |
| Ch09 | Di | NED | 321.931 | 321.931 | 322 |
| Ch09 | Di | NED | 324.249 | 325.308 | 324 |
| Ch09 | Di | NED | 325.763 | 326.378 | 326 |
| Ch09 | Di | NED | 327.876 | 328.768 | 328 |
| Ch09 | Di | NED | 330.907 | 330.907 | 330 |
| Ch09 | Di | NED | 336.354 | 336.354 | 336 |
| Ch09 | Di | NED | 343.219 | 343.219 | 342 |
| Ch09 | Di | NED | 350.68 | 351.263 | 350 |
| Ch09 | Di | NED | 360.35 | 360.35 | 360 |
| Ch09 | Di | NED | 362.199 | 362.199 | 362 |
| Ch43 | Di | 6-FAM | 151.823 | 151.823 | 150 |
| Ch43 | Di | 6-FAM | 153.43 | 153.729 | 152 |
| Ch43 | Di | 6-FAM | 154.856 | 155.278 | 154 |
| Ch43 | Di | 6-FAM | 160.775 | 161.637 | 160 |
| Ch43 | Di | 6-FAM | 162.891 | 162.891 | 162 |
| Ch43 | Di | 6-FAM | 164.805 | 165.656 | 164 |
| Ch43 | Di | 6-FAM | 166.256 | 167.111 | 166 |
| Ch43 | Di | 6-FAM | 170.591 | 170.591 | 170 |
| Ch43 | Di | 6-FAM | 171.588 | 172.586 | 172 |
| Ch43 | Di | 6-FAM | 173.377 | 173.377 | 174 |
| Ch45 | Tetra | 6-FAM | 111.826 | 111.895 | 112 |
| Ch45 | Tetra | 6-FAM | 115.771 | 116.164 | 116 |
| Ch45 | Tetra | 6-FAM | 119.762 | 120.65 | 120 |
| Ch45 | Tetra | 6-FAM | 124.117 | 124.562 | 124 |
| Ch45 | Tetra | 6-FAM | 128.22 | 128.712 | 128 |
| Ch54 | Di | PET | 231.501 | 231.501 | 231 |
| Ch54 | Di | PET | 237.401 | 238.06 | 237 |
| Ch54 | Di | PET | 238.763 | 239.784 | 239 |
| Ch54 | Di | PET | 241.613 | 242.186 | 241 |
| Ch54 | Di | PET | 243.739 | 243.974 | 243 |
| Ch54 | Di | PET | 245.89 | 245.995 | 245 |
| Ch54 | Di | PET | 251.856 | 252.137 | 251 |
| Ch54 | Di | PET | 256.622 | 256.622 | 257 |
| Ch54 | Di | PET | 266.675 | 266.675 | 267 |
| Ch54 | Di | PET | 268.149 | 268.149 | 269 |
| Ch54 | Di | PET | 272.056 | 272.056 | 271 |
| Ch54 | Di | PET | 278.14 | 278.608 | 277 |
| Ch54 | Di | PET | 280.292 | 280.44 | 279 |
| Ch54 | Di | PET | 282.22 | 282.661 | 281 |
| Ch54 | Di | PET | 284.579 | 284.584 | 285 |
| Ch54 | Di | PET | 292.561 | 292.808 | 293 |
| Ch60 | Tri | PET | 124.598 | 124.947 | 124 |
| Ch60 | Tri | PET | 145.578 | 145.789 | 145 |
| Ch60 | Tri | PET | 148.622 | 148.954 | 148 |
| Ch60 | Tri | PET | 151.72 | 152.119 | 151 |
| Ch63 | Tri | 6-FAM | 237.663 | 237.934 | 237 |
| Ch63 | Tri | 6-FAM | 240.702 | 240.844 | 240 |
| Ch63 | Tri | 6-FAM | 243.469 | 243.756 | 243 |
| Ch63 | Tri | 6-FAM | 246.332 | 246.871 | 246 |
| Ch63 | Tri | 6-FAM | 249.25 | 249.597 | 249 |
| Ch63 | Tri | 6-FAM | 253.307 | 253.307 | 253 |

**Table S5** Microsatellite locus information for *Chlainomonas* sp. Locus name, repeat motif, forward oligo fluorochrome, and poolplex information.

| **Locus** | **Motif** | **Dye** | **Poolplex** |
| --- | --- | --- | --- |
| Ch08 | (AG)_8_ | 6-FAM | 1 |
| Ch09 | (AC)_13_ | NED | 1 |
| Ch43 | (AG)_20_ | 6-FAM | Simplex |
| Ch45 | (AGGC)_7_ | 6-FAM | 2 |
| Ch54 | (AC)_11_ | PET | 1 |
| Ch60 | (AGC)_8_ | PET | Simplex |
| Ch63 | (AGC)_8_ | 6-FAM | 2 |

**Table S6** Null allele proportions at each site and time point by locus as calculated by maximum likelihood in ML-NullFreq (Kalinowski & Taper, 2006 *Conserv. Genet.* 7, 991–995 <https://doi.org/10.1007/s10592-006-9134-9>).

| **Locus** | **2022-June-02** | **2022-June-23** | **2022-June-30** | **2022-July-07** | **2025-June-05** | **2022** | **All Samples** |
| --- | --- | --- | --- | --- | --- | --- | --- |
| Ch08 | 0.37 | 0.138 | 0.21 | 0 | 0.727 | 0.414 | 0.499 |
| Ch09 | 0.36 | 0.377 | 0.387 | 0 | 0.402 | 0.382 | 0.39 |
| Ch43 | 0.223 | 0.236 | 0 | 0.574 | 0.469 | 0.281 | 0.358 |
| Ch45 | 0.341 | 0.523 | 0.431 | 0.571 | 0.256 | 0.405 | 0.362 |
| Ch54 | 0.141 | 0.162 | 0.278 | 0.176 | 0.052 | 0.208 | 0.17 |
| Ch60 | 0.554 | 0.776 | 0.353 | 0 | 0.35 | 0.549 | 0.489 |
| Ch63 | 0.322 | 0.483 | 0.256 | 0.315 | 0.332 | 0.343 | 0.341 |

**Table S7** Studies used to compare population *Pareto β* to *Chlainomonas sp.* Species, the species sampled; taxon, the taxonomic group of the species; reference, the publication source of the data; data reference, the specific source of the data if different from the published paper; life cycle type, whether the species has a haploid, diploid, or haploid-diploid life cycle; sample type, whether the microalgae were sampled from clonal cultures or single cells from wild blooms; bloom source, where cells of the given microalga originate from to start bloom; location, the place samples were gathered from in the study.

| **Species** | **Taxon** | **Reference** | **Image reference** | **Life cycle type** | **Sample type** | **Bloom source** | **Location** |
| --- | --- | --- | --- | --- | --- | --- | --- |
| *Alexandrium fundyense* | Dinoflagellate | Erdner et al. (1) | <https://www.marinespecies.org/aphia.php?p=image&pic=40247> | Haploid | Clonal culture | Sediment cysts | Gulf of Maine |
| *Alexandrium fundyense* | Dinoflagellate | Richlen et al. (2) | <https://www.marinespecies.org/aphia.php?p=image&pic=40247> | Haploid | Clonal culture | Sediment cysts | Cape Cod Pond |
| *Alexandrium minutum* | Dinoflagellate | Dia et al. (3) | Philippe Garcelon [https://www.flickr.com/photos/philgar/51841087009/in/photostream](https://www.flickr.com/photos/philgar/51841087009/in/photostream/) | Haploid | Clonal culture | Sediment cysts | Estuaries in Brittany, France |
| *Alexandrium ostenfeldii* | Dinoflagellate | Jerney et al. (4) | <https://www.marinespecies.org/photogallery.php?album=1033&pic=40248> | Haploid | Clonal culture | Sediment cysts | Baltic Sea |
| *Asterionella formosa* | Diatom | van den Wyngaert et al. (5) | [https://en.wikipedia.org/wiki/Asterionella_formosa#/media/File:Asterionella_formosa_mosbo6.jpg](https://en.wikipedia.org/wiki/Asterionella_formosa) | Diploid | Clonal culture | Cells in water column | Lakes in Switzerland and the Netherlands |
| *Gephyrocapsa (Emiliana) huxleyi* | Haptophyte | Krueger-Hadfield et al. (6) | [https://en.wikipedia.org/wiki/Gephyrocapsa_huxleyi#/media/File:Emiliania_huxleyi.jpg](https://en.wikipedia.org/wiki/Gephyrocapsa_huxleyi) | Haploid-diploid | Clonal culture | Cells in water column | North Sea |
| *Gambierdiscus caribaeus* | Dinoflagellate | Sassenhagen et al. (7) | <https://www.si.edu/es/object/gambierdiscus-sp:nmnhbotany_11257726> | Haploid | Clonal culture | Epiphytic cells on macroalgae | Florida Keys and US Virgin Islands |
| *Gonyostomum semen* | Raphidophyte | Rengefors et al. (8) | [https://en.wikipedia.org/wiki/Gonyostomum_semen#/media/File:Gonyostomum-cells.JPG](https://en.wikipedia.org/wiki/Gonyostomum_semen) | Haploid-diploid | Clonal culture | Sediment cysts | Scandinavia |
| *Gonyostomum semen* | Raphidophyte | Gollnisch et al. (9) | [https://en.wikipedia.org/wiki/Gonyostomum_semen#/media/File:Gonyostomum-cells.JPG](https://en.wikipedia.org/wiki/Gonyostomum_semen) | Haploid-diploid | Single cell | Sediment cysts | Lakes across Europe and the USA |
| *Prorocentrum donghaiense* | Dinoflagellate | Huang et al. (10) | <https://aquariumstoredepot.com/blogs/news/how-to-get-rid-of-dinoflagellates> | Haploid | Clonal culture | Cells in water column | East China Sea |
| *Pseudo-Nitzschia multiseries* | Diatom | Evans et al. (11) | [www.marinespecies.org/photogallery.php?pic=39697](http://www.marinespecies.org/photogallery.php?pic=39697) | Diploid | Clonal culture | Sediment cysts | Harbors around Atlantic Canada |
| *Pseudo-Nitzschia multistriata* | Diatom | Tesson et al. (12) | [www.marinespecies.org/photogallery.php?pic=39697](http://www.marinespecies.org/photogallery.php?pic=39697) | Diploid | Clonal culture | Sediment cysts | Gulf of Naples |
| *Pseudo-Nitzschia multistriata* | Diatom | Ruggiero et al. (13) | [www.marinespecies.org/photogallery.php?pic=39697](http://www.marinespecies.org/photogallery.php?pic=39697) | Diploid | Clonal culture | Sediment cysts | Gulf of Naples |
| *Pseudo-Nitzschia pungens* | Diatom | Adams et al. (14) | [www.marinespecies.org/photogallery.php?pic=39697](http://www.marinespecies.org/photogallery.php?pic=39697) | Diploid | Clonal culture | Sediment cysts | Juan de Fuca eddy, English Channel, and the North Sea |
| *Symbiodinium fitti* | Dinoflagellate | Baums et al. (15) | [https://en.wikipedia.org/wiki/Symbiodinium#/media/File:Symbiodinium.png](https://en.wikipedia.org/wiki/Symbiodinium) | Haploid | Clonal culture | Non bloom forming species | Caribbean Sea |

**Table S8a** Single locus values for expected heterozygosity (*H_E_*) at each patch and sampling date.

| **Population** | **Ch08** | **Ch09** | **Ch43** | **Ch45** | **Ch54** | **Ch60** | **Ch63** |
| --- | --- | --- | --- | --- | --- | --- | --- |
| 2022 | 0.64975 | 0.907518 | 0.1275 | 0.586356 | 0.778931 | 0.584548 | 0.658472 |
| 2022-June-02 | 0.701111 | 0.906746 | 0.09212 | 0.511898 | 0.704164 | 0.600551 | 0.674598 |
| 2022-June-23 | 0.489796 | 0.826531 | 0.244898 | 0.722222 | 0.714286 | 0.666667 | 0.5 |
| 2022-June-30 | 0.495868 | 0.780992 | 0.203704 | 0.567901 | 0.644628 | 0.375 | 0.51 |
| 2025-June-05 | 0.624444 | 0.866825 | 0.145429 | 0.57076 | 0.798889 | 0.615339 | 0.558264 |

**Table S8b** Single locus values for observed heterozygosity (*H_O_*) at each patch and sampling date.

| **Population** | **Ch08** | **Ch09** | **Ch43** | **Ch45** | **Ch54** | **Ch60** | **Ch63** |
| --- | --- | --- | --- | --- | --- | --- | --- |
| 2022 | 0.27451 | 0.241935 | 0.066667 | 0.137931 | 0.453125 | 0.061224 | 0.233333 |
| 2022-June-02 | 0.333333 | 0.285714 | 0.047619 | 0.146341 | 0.534884 | 0.060606 | 0.268293 |
| 2022-June-23 | 0.285714 | 0.142857 | 0 | 0 | 0.428571 | 0 | 0 |
| 2022-June-30 | 0.181818 | 0.090909 | 0.222222 | 0.222222 | 0.181818 | 0.1 | 0.3 |
| 2025-June-05 | 0.066667 | 0.206897 | 0.052632 | 0.193548 | 0.733333 | 0.172414 | 0.172414 |

**Table S8c** Single locus values for the inbreeding coefficient (*F_IS_*) at each patch and sampling date.

| **Population** | **Ch08** | **Ch09** | **Ch43** | **Ch45** | **Ch54** | **Ch60** | **Ch63** |
| --- | --- | --- | --- | --- | --- | --- | --- |
| 2022 | 0.577515 | 0.73341 | 0.477124 | 0.764766 | 0.418273 | 0.895262 | 0.645644 |
| 2022-June-02 | 0.524564 | 0.684902 | 0.483077 | 0.71412 | 0.240399 | 0.899083 | 0.602293 |
| 2022-June-23 | 0.416667 | 0.82716 | 1 | 1 | 0.4 | 1 | 1 |
| 2022-June-30 | 0.633333 | 0.883598 | -0.09091 | 0.608696 | 0.717949 | 0.733333 | 0.411765 |
| 2025-June-05 | 0.893238 | 0.761317 | 0.638095 | 0.660893 | 0.082058 | 0.719807 | 0.691161 |

**Table S9a** Posterior probabilities of rates of clonality, selfing, and outcrossing using changes of genotype frequencies between temporal samples (CEMP) gathered from Bagley Lake on 2 June 2022 and 23 June 2022. Outcrossing rate is calculated as 1 - clonal rate - selfing rate.

| **Mutation rate** | **Rate of clonality** | **Selfing rate** | | **Outcrossing rate** | **Posterior probability** | |
| --- | --- | --- | --- | --- | --- | --- |
| 0.001 | 0 | 0 | 1 | | | 3.20E-20 |
| 0.001 | 0.01 | 0 | 0.99 | | | 3.21E-19 |
| 0.001 | 0.02 | 0 | 0.98 | | | 2.20E-18 |
| 0.001 | 0.03 | 0 | 0.97 | | | 1.18E-17 |
| 0.001 | 0.04 | 0 | 0.96 | | | 5.27E-17 |
| 0.001 | 0.05 | 0 | 0.95 | | | 2.06E-16 |
| 0.001 | 0.06 | 0 | 0.94 | | | 7.26E-16 |
| 0.001 | 0.07 | 0 | 0.93 | | | 2.34E-15 |
| 0.001 | 0.08 | 0 | 0.92 | | | 6.98E-15 |
| 0.001 | 0.09 | 0 | 0.91 | | | 1.96E-14 |
| 0.001 | 0.1 | 0 | 0.9 | | | 5.19E-14 |
| 0.001 | 0.11 | 0 | 0.89 | | | 1.31E-13 |
| 0.001 | 0.12 | 0 | 0.88 | | | 3.17E-13 |
| 0.001 | 0.13 | 0 | 0.87 | | | 7.37E-13 |
| 0.001 | 0.14 | 0 | 0.86 | | | 1.65E-12 |
| 0.001 | 0.15 | 0 | 0.85 | | | 3.60E-12 |
| 0.001 | 0.16 | 0 | 0.84 | | | 7.59E-12 |
| 0.001 | 0.17 | 0 | 0.83 | | | 1.56E-11 |
| 0.001 | 0.18 | 0 | 0.82 | | | 3.13E-11 |
| 0.001 | 0.19 | 0 | 0.81 | | | 6.13E-11 |
| 0.001 | 0.2 | 0 | 0.8 | | | 1.17E-10 |
| 0.001 | 0.21 | 0 | 0.79 | | | 2.21E-10 |
| 0.001 | 0.22 | 0 | 0.78 | | | 4.07E-10 |
| 0.001 | 0.23 | 0 | 0.77 | | | 7.37E-10 |
| 0.001 | 0.24 | 0 | 0.76 | | | 1.31E-09 |
| 0.001 | 0.25 | 0 | 0.75 | | | 2.30E-09 |
| 0.001 | 0.26 | 0 | 0.74 | | | 3.97E-09 |
| 0.001 | 0.27 | 0 | 0.73 | | | 6.76E-09 |
| 0.001 | 0.28 | 0 | 0.72 | | | 1.14E-08 |
| 0.001 | 0.29 | 0 | 0.71 | | | 1.88E-08 |
| 0.001 | 0.3 | 0 | 0.7 | | | 3.08E-08 |
| 0.001 | 0.31 | 0 | 0.69 | | | 4.98E-08 |
| 0.001 | 0.32 | 0 | 0.68 | | | 7.97E-08 |
| 0.001 | 0.33 | 0 | 0.67 | | | 1.26E-07 |
| 0.001 | 0.34 | 0 | 0.66 | | | 1.98E-07 |
| 0.001 | 0.35 | 0 | 0.65 | | | 3.06E-07 |
| 0.001 | 0.36 | 0 | 0.64 | | | 4.70E-07 |
| 0.001 | 1 | 0 | 0 | | | 5.10E-07 |
| 0.001 | 0.37 | 0 | 0.63 | | | 7.14E-07 |
| 0.001 | 0.38 | 0 | 0.62 | | | 1.08E-06 |
| 0.001 | 0.39 | 0 | 0.61 | | | 1.61E-06 |
| 0.001 | 0.4 | 0 | 0.6 | | | 2.38E-06 |
| 0.001 | 0.41 | 0 | 0.59 | | | 3.49E-06 |
| 0.001 | 0.42 | 0 | 0.58 | | | 5.09E-06 |
| 0.001 | 0.43 | 0 | 0.57 | | | 7.35E-06 |
| 0.001 | 0.44 | 0 | 0.56 | | | 1.05E-05 |
| 0.001 | 0.45 | 0 | 0.55 | | | 1.50E-05 |
| 0.001 | 0.46 | 0 | 0.54 | | | 2.12E-05 |
| 0.001 | 0.47 | 0 | 0.53 | | | 2.97E-05 |
| 0.001 | 0.48 | 0 | 0.52 | | | 4.13E-05 |
| 0.001 | 0.49 | 0 | 0.51 | | | 5.71E-05 |
| 0.001 | 0.5 | 0 | 0.5 | | | 7.84E-05 |
| 0.001 | 0.51 | 0 | 0.49 | | | 0.000106891 |
| 0.001 | 0.99 | 0 | 0.01 | | | 0.000116675 |
| 0.001 | 0.52 | 0 | 0.48 | | | 0.000144773 |
| 0.001 | 0.53 | 0 | 0.47 | | | 0.000194784 |
| 0.001 | 0.54 | 0 | 0.46 | | | 0.000260357 |
| 0.001 | 0.55 | 0 | 0.45 | | | 0.000345747 |
| 0.001 | 0.56 | 0 | 0.44 | | | 0.000456185 |
| 0.001 | 0.57 | 0 | 0.43 | | | 0.000598041 |
| 0.001 | 0.58 | 0 | 0.42 | | | 0.000779 |
| 0.001 | 0.98 | 0 | 0.02 | | | 0.000871058 |
| 0.001 | 0.59 | 0 | 0.41 | | | 0.001008242 |
| 0.001 | 0.6 | 0 | 0.4 | | | 0.001296618 |
| 0.001 | 0.61 | 0 | 0.39 | | | 0.00165681 |
| 0.001 | 0.62 | 0 | 0.38 | | | 0.002103461 |
| 0.001 | 0.63 | 0 | 0.37 | | | 0.002653256 |
| 0.001 | 0.97 | 0 | 0.03 | | | 0.002853861 |
| 0.001 | 0.64 | 0 | 0.36 | | | 0.003324928 |
| 0.001 | 0.65 | 0 | 0.35 | | | 0.004139157 |
| 0.001 | 0.66 | 0 | 0.34 | | | 0.005118342 |
| 0.001 | 0.67 | 0 | 0.33 | | | 0.006286186 |
| 0.001 | 0.96 | 0 | 0.04 | | | 0.006380694 |
| 0.001 | 0.68 | 0 | 0.32 | | | 0.007667077 |
| 0.001 | 0.69 | 0 | 0.31 | | | 0.009285206 |
| 0.001 | 0.7 | 0 | 0.3 | | | 0.011163384 |
| 0.001 | 0.95 | 0 | 0.05 | | | 0.011424745 |
| 0.001 | 0.71 | 0 | 0.29 | | | 0.013321544 |
| 0.001 | 0.72 | 0 | 0.28 | | | 0.015774872 |
| 0.001 | 0.94 | 0 | 0.06 | | | 0.017681487 |
| 0.001 | 0.73 | 0 | 0.27 | | | 0.018531593 |
| 0.001 | 0.74 | 0 | 0.26 | | | 0.021590423 |
| 0.001 | 0.93 | 0 | 0.07 | | | 0.024677769 |
| 0.001 | 0.75 | 0 | 0.25 | | | 0.024937743 |
| 0.001 | 0.76 | 0 | 0.24 | | | 0.028544615 |
| 0.001 | 0.92 | 0 | 0.08 | | | 0.031879183 |
| 0.001 | 0.77 | 0 | 0.23 | | | 0.032363793 |
| 0.001 | 0.78 | 0 | 0.22 | | | 0.03632697 |
| 0.001 | 0.91 | 0 | 0.09 | | | 0.038775123 |
| 0.001 | 0.79 | 0 | 0.21 | | | 0.040342576 |
| 0.001 | 0.8 | 0 | 0.2 | | | 0.04429451 |
| 0.001 | 0.9 | 0 | 0.1 | | | 0.044935556 |
| 0.001 | 0.81 | 0 | 0.19 | | | 0.048042285 |
| 0.001 | 0.89 | 0 | 0.11 | | | 0.050041225 |
| 0.001 | 0.82 | 0 | 0.18 | | | 0.051423098 |
| 0.001 | 0.88 | 0 | 0.12 | | | 0.05389245 |
| 0.001 | 0.83 | 0 | 0.17 | | | 0.05425639 |
| 0.001 | 0.84 | 0 | 0.16 | | | 0.056351413 |
| 0.001 | 0.87 | 0 | 0.13 | | | 0.056402648 |
| 0.001 | 0.85 | 0 | 0.15 | | | 0.057518227 |
| 0.001 | 0.86 | 0 | 0.14 | | | 0.057582317 |

**Table S9b** Posterior probabilities of rates of clonality, selfing, and outcrossing using changes of genotype frequencies between temporal samples (CEMP) gathered from Bagley Lake on 2 June 2022 and 30 June 2022. Outcrossing rate is calculated as 1 - clonal rate - selfing rate.

| **Mutation rate** | **Rate of clonality** | **Selfing rate** | **Outcrossing rate** | **Posterior probability** |
| --- | --- | --- | --- | --- |
| 0.001 | 0 | 0 | 1 | 1.50E-20 |
| 0.001 | 0.01 | 0 | 0.99 | 8.98E-20 |
| 0.001 | 0.02 | 0 | 0.98 | 4.65E-19 |
| 0.001 | 0.03 | 0 | 0.97 | 2.15E-18 |
| 0.001 | 0.04 | 0 | 0.96 | 9.00E-18 |
| 0.001 | 0.05 | 0 | 0.95 | 3.48E-17 |
| 0.001 | 0.06 | 0 | 0.94 | 1.26E-16 |
| 0.001 | 0.07 | 0 | 0.93 | 4.26E-16 |
| 0.001 | 0.08 | 0 | 0.92 | 1.37E-15 |
| 0.001 | 1 | 0 | 0 | 1.74E-15 |
| 0.001 | 0.09 | 0 | 0.91 | 4.17E-15 |
| 0.001 | 0.1 | 0 | 0.9 | 1.22E-14 |
| 0.001 | 0.11 | 0 | 0.89 | 3.41E-14 |
| 0.001 | 0.12 | 0 | 0.88 | 9.21E-14 |
| 0.001 | 0.13 | 0 | 0.87 | 2.40E-13 |
| 0.001 | 0.14 | 0 | 0.86 | 6.05E-13 |
| 0.001 | 0.15 | 0 | 0.85 | 1.48E-12 |
| 0.001 | 0.16 | 0 | 0.84 | 3.52E-12 |
| 0.001 | 0.17 | 0 | 0.83 | 8.14E-12 |
| 0.001 | 0.18 | 0 | 0.82 | 1.84E-11 |
| 0.001 | 0.19 | 0 | 0.81 | 4.04E-11 |
| 0.001 | 0.2 | 0 | 0.8 | 8.71E-11 |
| 0.001 | 0.21 | 0 | 0.79 | 1.83E-10 |
| 0.001 | 0.99 | 0 | 0.01 | 3.00E-10 |
| 0.001 | 0.22 | 0 | 0.78 | 3.79E-10 |
| 0.001 | 0.23 | 0 | 0.77 | 7.66E-10 |
| 0.001 | 0.24 | 0 | 0.76 | 1.52E-09 |
| 0.001 | 0.25 | 0 | 0.75 | 2.96E-09 |
| 0.001 | 0.26 | 0 | 0.74 | 5.67E-09 |
| 0.001 | 0.27 | 0 | 0.73 | 1.07E-08 |
| 0.001 | 0.28 | 0 | 0.72 | 1.98E-08 |
| 0.001 | 0.98 | 0 | 0.02 | 3.48E-08 |
| 0.001 | 0.29 | 0 | 0.71 | 3.61E-08 |
| 0.001 | 0.3 | 0 | 0.7 | 6.48E-08 |
| 0.001 | 0.31 | 0 | 0.69 | 1.15E-07 |
| 0.001 | 0.32 | 0 | 0.68 | 2.00E-07 |
| 0.001 | 0.33 | 0 | 0.67 | 3.44E-07 |
| 0.001 | 0.34 | 0 | 0.66 | 5.83E-07 |
| 0.001 | 0.97 | 0 | 0.03 | 6.59E-07 |
| 0.001 | 0.35 | 0 | 0.65 | 9.75E-07 |
| 0.001 | 0.36 | 0 | 0.64 | 1.61E-06 |
| 0.001 | 0.37 | 0 | 0.63 | 2.63E-06 |
| 0.001 | 0.38 | 0 | 0.62 | 4.23E-06 |
| 0.001 | 0.96 | 0 | 0.04 | 5.35E-06 |
| 0.001 | 0.39 | 0 | 0.61 | 6.73E-06 |
| 0.001 | 0.4 | 0 | 0.6 | 1.06E-05 |
| 0.001 | 0.41 | 0 | 0.59 | 1.64E-05 |
| 0.001 | 0.42 | 0 | 0.58 | 2.52E-05 |
| 0.001 | 0.95 | 0 | 0.05 | 2.64E-05 |
| 0.001 | 0.43 | 0 | 0.57 | 3.83E-05 |
| 0.001 | 0.44 | 0 | 0.56 | 5.74E-05 |
| 0.001 | 0.45 | 0 | 0.55 | 8.52E-05 |
| 0.001 | 0.94 | 0 | 0.06 | 9.44E-05 |
| 0.001 | 0.46 | 0 | 0.54 | 0.000124938 |
| 0.001 | 0.47 | 0 | 0.53 | 0.000181283 |
| 0.001 | 0.48 | 0 | 0.52 | 0.000260166 |
| 0.001 | 0.93 | 0 | 0.07 | 0.000267464 |
| 0.001 | 0.49 | 0 | 0.51 | 0.000369322 |
| 0.001 | 0.5 | 0 | 0.5 | 0.000518608 |
| 0.001 | 0.92 | 0 | 0.08 | 0.000637111 |
| 0.001 | 0.51 | 0 | 0.49 | 0.000720387 |
| 0.001 | 0.52 | 0 | 0.48 | 0.000989899 |
| 0.001 | 0.91 | 0 | 0.09 | 0.001325814 |
| 0.001 | 0.53 | 0 | 0.47 | 0.00134559 |
| 0.001 | 0.54 | 0 | 0.46 | 0.001809352 |
| 0.001 | 0.55 | 0 | 0.45 | 0.002406606 |
| 0.001 | 0.9 | 0 | 0.1 | 0.002475149 |
| 0.001 | 0.56 | 0 | 0.44 | 0.003166173 |
| 0.001 | 0.57 | 0 | 0.43 | 0.004119834 |
| 0.001 | 0.89 | 0 | 0.11 | 0.004225777 |
| 0.001 | 0.58 | 0 | 0.42 | 0.005301504 |
| 0.001 | 0.88 | 0 | 0.12 | 0.006692947 |
| 0.001 | 0.59 | 0 | 0.41 | 0.006745942 |
| 0.001 | 0.6 | 0 | 0.4 | 0.008486927 |
| 0.001 | 0.87 | 0 | 0.13 | 0.009942548 |
| 0.001 | 0.61 | 0 | 0.39 | 0.010554866 |
| 0.001 | 0.62 | 0 | 0.38 | 0.012973831 |
| 0.001 | 0.86 | 0 | 0.14 | 0.013972587 |
| 0.001 | 0.63 | 0 | 0.37 | 0.015758106 |
| 0.001 | 0.85 | 0 | 0.15 | 0.018703742 |
| 0.001 | 0.64 | 0 | 0.36 | 0.018908373 |
| 0.001 | 0.65 | 0 | 0.35 | 0.022407775 |
| 0.001 | 0.84 | 0 | 0.16 | 0.02398067 |
| 0.001 | 0.66 | 0 | 0.34 | 0.026218174 |
| 0.001 | 0.83 | 0 | 0.17 | 0.029583719 |
| 0.001 | 0.67 | 0 | 0.33 | 0.030277011 |
| 0.001 | 0.68 | 0 | 0.32 | 0.034495262 |
| 0.001 | 0.82 | 0 | 0.18 | 0.035248906 |
| 0.001 | 0.69 | 0 | 0.31 | 0.038756988 |
| 0.001 | 0.81 | 0 | 0.19 | 0.040692913 |
| 0.001 | 0.7 | 0 | 0.3 | 0.042920988 |
| 0.001 | 0.8 | 0 | 0.2 | 0.045639411 |
| 0.001 | 0.71 | 0 | 0.29 | 0.04682496 |
| 0.001 | 0.79 | 0 | 0.21 | 0.049843259 |
| 0.001 | 0.72 | 0 | 0.28 | 0.050292423 |
| 0.001 | 0.78 | 0 | 0.22 | 0.053109871 |
| 0.001 | 0.73 | 0 | 0.27 | 0.053142399 |
| 0.001 | 0.74 | 0 | 0.26 | 0.055201524 |
| 0.001 | 0.77 | 0 | 0.23 | 0.055308027 |
| 0.001 | 0.75 | 0 | 0.25 | 0.056317887 |
| 0.001 | 0.76 | 0 | 0.24 | 0.056375511 |

**Table S9c** Posterior probabilities of rates of clonality, selfing, and outcrossing using changes of genotype frequencies between temporal samples (CEMP) gathered from Bagley Lake on 2 June 2022 and 5 June 2025. Outcrossing rate is calculated as 1 - clonal rate - selfing rate.

| **Mutation rate** | | **Rate of clonality** | **Selfing rate** | | **Outcrossing rate** | **Posterior probability** |
| --- | --- | --- | --- | --- | --- | --- |
| 0.001 | 0 | | | 0 | 1 | 1.19E-85 |
| 0.001 | 0.01 | | | 0 | 0.99 | 7.22E-82 |
| 0.001 | 0.02 | | | 0 | 0.98 | 1.29E-78 |
| 0.001 | 0.03 | | | 0 | 0.97 | 9.97E-76 |
| 0.001 | 0.04 | | | 0 | 0.96 | 4.18E-73 |
| 0.001 | 0.05 | | | 0 | 0.95 | 1.08E-70 |
| 0.001 | 0.06 | | | 0 | 0.94 | 1.90E-68 |
| 0.001 | 0.07 | | | 0 | 0.93 | 2.42E-66 |
| 0.001 | 0.08 | | | 0 | 0.92 | 2.33E-64 |
| 0.001 | 0.09 | | | 0 | 0.91 | 1.78E-62 |
| 0.001 | 0.1 | | | 0 | 0.9 | 1.10E-60 |
| 0.001 | 0.11 | | | 0 | 0.89 | 5.64E-59 |
| 0.001 | 0.12 | | | 0 | 0.88 | 2.45E-57 |
| 0.001 | 0.13 | | | 0 | 0.87 | 9.20E-56 |
| 0.001 | 0.14 | | | 0 | 0.86 | 3.01E-54 |
| 0.001 | 0.15 | | | 0 | 0.85 | 8.72E-53 |
| 0.001 | 0.16 | | | 0 | 0.84 | 2.25E-51 |
| 0.001 | 0.17 | | | 0 | 0.83 | 5.24E-50 |
| 0.001 | 0.18 | | | 0 | 0.82 | 1.10E-48 |
| 0.001 | 0.19 | | | 0 | 0.81 | 2.13E-47 |
| 0.001 | 0.2 | | | 0 | 0.8 | 3.76E-46 |
| 0.001 | 0.21 | | | 0 | 0.79 | 6.15E-45 |
| 0.001 | 0.22 | | | 0 | 0.78 | 9.32E-44 |
| 0.001 | 0.23 | | | 0 | 0.77 | 1.32E-42 |
| 0.001 | 0.24 | | | 0 | 0.76 | 1.74E-41 |
| 0.001 | 0.25 | | | 0 | 0.75 | 2.15E-40 |
| 0.001 | 0.26 | | | 0 | 0.74 | 2.52E-39 |
| 0.001 | 0.27 | | | 0 | 0.73 | 2.77E-38 |
| 0.001 | 0.28 | | | 0 | 0.72 | 2.90E-37 |
| 0.001 | 0.29 | | | 0 | 0.71 | 2.87E-36 |
| 0.001 | 0.3 | | | 0 | 0.7 | 2.70E-35 |
| 0.001 | 0.31 | | | 0 | 0.69 | 2.43E-34 |
| 0.001 | 0.32 | | | 0 | 0.68 | 2.08E-33 |
| 0.001 | 0.33 | | | 0 | 0.67 | 1.71E-32 |
| 0.001 | 0.34 | | | 0 | 0.66 | 1.35E-31 |
| 0.001 | 0.35 | | | 0 | 0.65 | 1.02E-30 |
| 0.001 | 0.36 | | | 0 | 0.64 | 7.38E-30 |
| 0.001 | 0.37 | | | 0 | 0.63 | 5.15E-29 |
| 0.001 | 0.38 | | | 0 | 0.62 | 3.47E-28 |
| 0.001 | 0.39 | | | 0 | 0.61 | 2.25E-27 |
| 0.001 | 0.4 | | | 0 | 0.6 | 1.41E-26 |
| 0.001 | 0.41 | | | 0 | 0.59 | 8.57E-26 |
| 0.001 | 0.42 | | | 0 | 0.58 | 5.03E-25 |
| 0.001 | 0.43 | | | 0 | 0.57 | 2.86E-24 |
| 0.001 | 0.44 | | | 0 | 0.56 | 1.58E-23 |
| 0.001 | 0.45 | | | 0 | 0.55 | 8.42E-23 |
| 0.001 | 0.46 | | | 0 | 0.54 | 4.37E-22 |
| 0.001 | 0.47 | | | 0 | 0.53 | 2.20E-21 |
| 0.001 | 0.48 | | | 0 | 0.52 | 1.08E-20 |
| 0.001 | 0.49 | | | 0 | 0.51 | 5.15E-20 |
| 0.001 | 0.5 | | | 0 | 0.5 | 2.39E-19 |
| 0.001 | 0.51 | | | 0 | 0.49 | 1.08E-18 |
| 0.001 | 0.52 | | | 0 | 0.48 | 4.75E-18 |
| 0.001 | 0.53 | | | 0 | 0.47 | 2.04E-17 |
| 0.001 | 0.54 | | | 0 | 0.46 | 8.51E-17 |
| 0.001 | 0.55 | | | 0 | 0.45 | 3.47E-16 |
| 0.001 | 0.56 | | | 0 | 0.44 | 1.38E-15 |
| 0.001 | 0.57 | | | 0 | 0.43 | 5.37E-15 |
| 0.001 | 0.58 | | | 0 | 0.42 | 2.04E-14 |
| 0.001 | 0.59 | | | 0 | 0.41 | 7.54E-14 |
| 0.001 | 0.6 | | | 0 | 0.4 | 2.73E-13 |
| 0.001 | 0.61 | | | 0 | 0.39 | 9.63E-13 |
| 0.001 | 0.62 | | | 0 | 0.38 | 3.33E-12 |
| 0.001 | 0.63 | | | 0 | 0.37 | 1.12E-11 |
| 0.001 | 0.64 | | | 0 | 0.36 | 3.70E-11 |
| 0.001 | 0.65 | | | 0 | 0.35 | 1.19E-10 |
| 0.001 | 0.66 | | | 0 | 0.34 | 3.75E-10 |
| 0.001 | 0.67 | | | 0 | 0.33 | 1.15E-09 |
| 0.001 | 0.68 | | | 0 | 0.32 | 3.47E-09 |
| 0.001 | 0.69 | | | 0 | 0.31 | 1.02E-08 |
| 0.001 | 0.7 | | | 0 | 0.3 | 2.92E-08 |
| 0.001 | 0.71 | | | 0 | 0.29 | 8.18E-08 |
| 0.001 | 1 | | | 0 | 0 | 8.26E-08 |
| 0.001 | 0.72 | | | 0 | 0.28 | 2.24E-07 |
| 0.001 | 0.73 | | | 0 | 0.27 | 5.97E-07 |
| 0.001 | 0.74 | | | 0 | 0.26 | 1.55E-06 |
| 0.001 | 0.75 | | | 0 | 0.25 | 3.94E-06 |
| 0.001 | 0.76 | | | 0 | 0.24 | 9.75E-06 |
| 0.001 | 0.77 | | | 0 | 0.23 | 2.35E-05 |
| 0.001 | 0.78 | | | 0 | 0.22 | 5.50E-05 |
| 0.001 | 0.79 | | | 0 | 0.21 | 0.000125056 |
| 0.001 | 0.8 | | | 0 | 0.2 | 0.000276153 |
| 0.001 | 0.99 | | | 0 | 0.01 | 0.000364872 |
| 0.001 | 0.81 | | | 0 | 0.19 | 0.000591016 |
| 0.001 | 0.82 | | | 0 | 0.18 | 0.001223856 |
| 0.001 | 0.83 | | | 0 | 0.17 | 0.002447307 |
| 0.001 | 0.84 | | | 0 | 0.16 | 0.004714744 |
| 0.001 | 0.98 | | | 0 | 0.02 | 0.005959123 |
| 0.001 | 0.85 | | | 0 | 0.15 | 0.008726142 |
| 0.001 | 0.86 | | | 0 | 0.14 | 0.015463595 |
| 0.001 | 0.87 | | | 0 | 0.13 | 0.02612931 |
| 0.001 | 0.97 | | | 0 | 0.03 | 0.026638653 |
| 0.001 | 0.88 | | | 0 | 0.12 | 0.041884893 |
| 0.001 | 0.89 | | | 0 | 0.11 | 0.06328774 |
| 0.001 | 0.96 | | | 0 | 0.04 | 0.064073576 |
| 0.001 | 0.9 | | | 0 | 0.1 | 0.089406715 |
| 0.001 | 0.95 | | | 0 | 0.05 | 0.10639862 |
| 0.001 | 0.91 | | | 0 | 0.09 | 0.116840793 |
| 0.001 | 0.94 | | | 0 | 0.06 | 0.13763633 |
| 0.001 | 0.92 | | | 0 | 0.08 | 0.139261621 |
| 0.001 | 0.93 | | | 0 | 0.07 | 0.148455167 |

**Table S9d** Posterior probabilities of rates of clonality, selfing, and outcrossing using changes of genotype frequencies between temporal samples (CEMP) gathered from Bagley Lake on 23 June 2022 and 30 June 2022. Outcrossing rate is calculated as 1 - clonal rate - selfing rate.

| **Mutation rate** | **Rate of clonality** | **Selfing rate** | **Outcrossing rate** | **Posterior probability** |
| --- | --- | --- | --- | --- |
| 0.001 | 0 | 0 | 1 | 2.33E-20 |
| 0.001 | 0.01 | 0 | 0.99 | 8.95E-20 |
| 0.001 | 0.02 | 0 | 0.98 | 3.23E-19 |
| 0.001 | 0.03 | 0 | 0.97 | 1.10E-18 |
| 0.001 | 0.04 | 0 | 0.96 | 3.53E-18 |
| 0.001 | 0.05 | 0 | 0.95 | 1.08E-17 |
| 0.001 | 0.06 | 0 | 0.94 | 3.17E-17 |
| 0.001 | 0.07 | 0 | 0.93 | 8.93E-17 |
| 0.001 | 0.08 | 0 | 0.92 | 2.42E-16 |
| 0.001 | 0.09 | 0 | 0.91 | 6.33E-16 |
| 0.001 | 0.1 | 0 | 0.9 | 1.60E-15 |
| 0.001 | 0.11 | 0 | 0.89 | 3.94E-15 |
| 0.001 | 0.12 | 0 | 0.88 | 9.40E-15 |
| 0.001 | 0.13 | 0 | 0.87 | 2.18E-14 |
| 0.001 | 0.14 | 0 | 0.86 | 4.95E-14 |
| 0.001 | 0.15 | 0 | 0.85 | 1.10E-13 |
| 0.001 | 0.16 | 0 | 0.84 | 2.37E-13 |
| 0.001 | 0.17 | 0 | 0.83 | 5.03E-13 |
| 0.001 | 0.18 | 0 | 0.82 | 1.05E-12 |
| 0.001 | 0.19 | 0 | 0.81 | 2.13E-12 |
| 0.001 | 0.2 | 0 | 0.8 | 4.26E-12 |
| 0.001 | 0.21 | 0 | 0.79 | 8.39E-12 |
| 0.001 | 0.22 | 0 | 0.78 | 1.62E-11 |
| 0.001 | 0.23 | 0 | 0.77 | 3.10E-11 |
| 0.001 | 0.24 | 0 | 0.76 | 5.81E-11 |
| 0.001 | 0.25 | 0 | 0.75 | 1.08E-10 |
| 0.001 | 0.26 | 0 | 0.74 | 1.96E-10 |
| 0.001 | 0.27 | 0 | 0.73 | 3.53E-10 |
| 0.001 | 0.28 | 0 | 0.72 | 6.28E-10 |
| 0.001 | 0.29 | 0 | 0.71 | 1.10E-09 |
| 0.001 | 0.3 | 0 | 0.7 | 1.91E-09 |
| 0.001 | 0.31 | 0 | 0.69 | 3.28E-09 |
| 0.001 | 0.32 | 0 | 0.68 | 5.57E-09 |
| 0.001 | 0.33 | 0 | 0.67 | 9.35E-09 |
| 0.001 | 0.34 | 0 | 0.66 | 1.55E-08 |
| 0.001 | 0.35 | 0 | 0.65 | 2.55E-08 |
| 0.001 | 0.36 | 0 | 0.64 | 4.16E-08 |
| 0.001 | 0.37 | 0 | 0.63 | 6.71E-08 |
| 0.001 | 0.38 | 0 | 0.62 | 1.07E-07 |
| 0.001 | 0.39 | 0 | 0.61 | 1.70E-07 |
| 0.001 | 0.4 | 0 | 0.6 | 2.67E-07 |
| 0.001 | 0.41 | 0 | 0.59 | 4.15E-07 |
| 0.001 | 1 | 0 | 0 | 4.86E-07 |
| 0.001 | 0.42 | 0 | 0.58 | 6.40E-07 |
| 0.001 | 0.43 | 0 | 0.57 | 9.80E-07 |
| 0.001 | 0.44 | 0 | 0.56 | 1.49E-06 |
| 0.001 | 0.45 | 0 | 0.55 | 2.24E-06 |
| 0.001 | 0.46 | 0 | 0.54 | 3.35E-06 |
| 0.001 | 0.47 | 0 | 0.53 | 4.97E-06 |
| 0.001 | 0.48 | 0 | 0.52 | 7.32E-06 |
| 0.001 | 0.49 | 0 | 0.51 | 1.07E-05 |
| 0.001 | 0.5 | 0 | 0.5 | 1.55E-05 |
| 0.001 | 0.51 | 0 | 0.49 | 2.23E-05 |
| 0.001 | 0.52 | 0 | 0.48 | 3.20E-05 |
| 0.001 | 0.53 | 0 | 0.47 | 4.54E-05 |
| 0.001 | 0.54 | 0 | 0.46 | 6.40E-05 |
| 0.001 | 0.55 | 0 | 0.45 | 8.97E-05 |
| 0.001 | 0.56 | 0 | 0.44 | 0.000124771 |
| 0.001 | 0.57 | 0 | 0.43 | 0.000172386 |
| 0.001 | 0.99 | 0 | 0.01 | 0.000194207 |
| 0.001 | 0.58 | 0 | 0.42 | 0.00023655 |
| 0.001 | 0.59 | 0 | 0.41 | 0.000322392 |
| 0.001 | 0.6 | 0 | 0.4 | 0.000436399 |
| 0.001 | 0.61 | 0 | 0.39 | 0.000586697 |
| 0.001 | 0.62 | 0 | 0.38 | 0.000783362 |
| 0.001 | 0.63 | 0 | 0.37 | 0.001038752 |
| 0.001 | 0.64 | 0 | 0.36 | 0.001367845 |
| 0.001 | 0.98 | 0 | 0.02 | 0.001462679 |
| 0.001 | 0.65 | 0 | 0.35 | 0.001788575 |
| 0.001 | 0.66 | 0 | 0.34 | 0.002322113 |
| 0.001 | 0.67 | 0 | 0.33 | 0.002993086 |
| 0.001 | 0.68 | 0 | 0.32 | 0.003829648 |
| 0.001 | 0.97 | 0 | 0.03 | 0.004694591 |
| 0.001 | 0.69 | 0 | 0.31 | 0.004863371 |
| 0.001 | 0.7 | 0 | 0.3 | 0.006128851 |
| 0.001 | 0.71 | 0 | 0.29 | 0.007662947 |
| 0.001 | 0.72 | 0 | 0.28 | 0.009503563 |
| 0.001 | 0.96 | 0 | 0.04 | 0.010198473 |
| 0.001 | 0.73 | 0 | 0.27 | 0.011687849 |
| 0.001 | 0.74 | 0 | 0.26 | 0.014249723 |
| 0.001 | 0.75 | 0 | 0.25 | 0.017216644 |
| 0.001 | 0.95 | 0 | 0.05 | 0.017679189 |
| 0.001 | 0.76 | 0 | 0.24 | 0.020605566 |
| 0.001 | 0.77 | 0 | 0.23 | 0.024418103 |
| 0.001 | 0.94 | 0 | 0.06 | 0.026437458 |
| 0.001 | 0.78 | 0 | 0.22 | 0.028634991 |
| 0.001 | 0.79 | 0 | 0.21 | 0.033210086 |
| 0.001 | 0.93 | 0 | 0.07 | 0.035606433 |
| 0.001 | 0.8 | 0 | 0.2 | 0.03806428 |
| 0.001 | 0.81 | 0 | 0.19 | 0.043079932 |
| 0.001 | 0.92 | 0 | 0.08 | 0.044344444 |
| 0.001 | 0.82 | 0 | 0.18 | 0.048096628 |
| 0.001 | 0.91 | 0 | 0.09 | 0.0519597 |
| 0.001 | 0.83 | 0 | 0.17 | 0.052909346 |
| 0.001 | 0.84 | 0 | 0.16 | 0.057270319 |
| 0.001 | 0.9 | 0 | 0.1 | 0.057970541 |
| 0.001 | 0.85 | 0 | 0.15 | 0.060896047 |
| 0.001 | 0.89 | 0 | 0.11 | 0.062115962 |
| 0.001 | 0.86 | 0 | 0.14 | 0.063480946 |
| 0.001 | 0.88 | 0 | 0.12 | 0.06433341 |
| 0.001 | 0.87 | 0 | 0.13 | 0.0647189 |

**Table S9e** Posterior probabilities of rates of clonality, selfing, and outcrossing using changes of genotype frequencies between temporal samples (CEMP) gathered from Bagley Lake on 23 June 2022 and 5 June 2025. Outcrossing rate is calculated as 1 - clonal rate - selfing rate.

| **Mutation rate** | **Rate of clonality** | **Selfing rate** | **Outcrossing rate** | **Posterior probability** |
| --- | --- | --- | --- | --- |
| 0.001 | 0 | 0 | 1 | 5.02E-44 |
| 0.001 | 0.01 | 0 | 0.99 | 1.24E-42 |
| 0.001 | 0.02 | 0 | 0.98 | 2.65E-41 |
| 0.001 | 0.03 | 0 | 0.97 | 4.91E-40 |
| 0.001 | 0.04 | 0 | 0.96 | 8.05E-39 |
| 0.001 | 0.05 | 0 | 0.95 | 1.18E-37 |
| 0.001 | 0.06 | 0 | 0.94 | 1.55E-36 |
| 0.001 | 0.07 | 0 | 0.93 | 1.86E-35 |
| 0.001 | 0.08 | 0 | 0.92 | 2.03E-34 |
| 0.001 | 0.09 | 0 | 0.91 | 2.05E-33 |
| 0.001 | 0.1 | 0 | 0.9 | 1.91E-32 |
| 0.001 | 0.11 | 0 | 0.89 | 1.65E-31 |
| 0.001 | 0.12 | 0 | 0.88 | 1.34E-30 |
| 0.001 | 0.13 | 0 | 0.87 | 1.02E-29 |
| 0.001 | 0.14 | 0 | 0.86 | 7.31E-29 |
| 0.001 | 0.15 | 0 | 0.85 | 4.95E-28 |
| 0.001 | 0.16 | 0 | 0.84 | 3.17E-27 |
| 0.001 | 0.17 | 0 | 0.83 | 1.94E-26 |
| 0.001 | 0.18 | 0 | 0.82 | 1.12E-25 |
| 0.001 | 0.19 | 0 | 0.81 | 6.24E-25 |
| 0.001 | 1 | 0 | 0 | 2.44E-24 |
| 0.001 | 0.2 | 0 | 0.8 | 3.31E-24 |
| 0.001 | 0.21 | 0 | 0.79 | 1.69E-23 |
| 0.001 | 0.22 | 0 | 0.78 | 8.26E-23 |
| 0.001 | 0.23 | 0 | 0.77 | 3.89E-22 |
| 0.001 | 0.24 | 0 | 0.76 | 1.77E-21 |
| 0.001 | 0.25 | 0 | 0.75 | 7.73E-21 |
| 0.001 | 0.26 | 0 | 0.74 | 3.27E-20 |
| 0.001 | 0.27 | 0 | 0.73 | 1.34E-19 |
| 0.001 | 0.28 | 0 | 0.72 | 5.32E-19 |
| 0.001 | 0.29 | 0 | 0.71 | 2.05E-18 |
| 0.001 | 0.3 | 0 | 0.7 | 7.65E-18 |
| 0.001 | 0.31 | 0 | 0.69 | 2.78E-17 |
| 0.001 | 0.32 | 0 | 0.68 | 9.80E-17 |
| 0.001 | 0.33 | 0 | 0.67 | 3.37E-16 |
| 0.001 | 0.34 | 0 | 0.66 | 1.13E-15 |
| 0.001 | 0.35 | 0 | 0.65 | 3.67E-15 |
| 0.001 | 0.36 | 0 | 0.64 | 1.17E-14 |
| 0.001 | 0.37 | 0 | 0.63 | 3.63E-14 |
| 0.001 | 0.38 | 0 | 0.62 | 1.10E-13 |
| 0.001 | 0.99 | 0 | 0.01 | 1.66E-13 |
| 0.001 | 0.39 | 0 | 0.61 | 3.26E-13 |
| 0.001 | 0.4 | 0 | 0.6 | 9.42E-13 |
| 0.001 | 0.41 | 0 | 0.59 | 2.67E-12 |
| 0.001 | 0.42 | 0 | 0.58 | 7.39E-12 |
| 0.001 | 0.43 | 0 | 0.57 | 2.00E-11 |
| 0.001 | 0.44 | 0 | 0.56 | 5.31E-11 |
| 0.001 | 0.45 | 0 | 0.55 | 1.38E-10 |
| 0.001 | 0.46 | 0 | 0.54 | 3.51E-10 |
| 0.001 | 0.98 | 0 | 0.02 | 5.64E-10 |
| 0.001 | 0.47 | 0 | 0.53 | 8.75E-10 |
| 0.001 | 0.48 | 0 | 0.52 | 2.14E-09 |
| 0.001 | 0.49 | 0 | 0.51 | 5.13E-09 |
| 0.001 | 0.5 | 0 | 0.5 | 1.20E-08 |
| 0.001 | 0.51 | 0 | 0.49 | 2.77E-08 |
| 0.001 | 0.52 | 0 | 0.48 | 6.26E-08 |
| 0.001 | 0.97 | 0 | 0.03 | 6.79E-08 |
| 0.001 | 0.53 | 0 | 0.47 | 1.39E-07 |
| 0.001 | 0.54 | 0 | 0.46 | 3.01E-07 |
| 0.001 | 0.55 | 0 | 0.45 | 6.42E-07 |
| 0.001 | 0.56 | 0 | 0.44 | 1.34E-06 |
| 0.001 | 0.96 | 0 | 0.04 | 1.81E-06 |
| 0.001 | 0.57 | 0 | 0.43 | 2.75E-06 |
| 0.001 | 0.58 | 0 | 0.42 | 5.53E-06 |
| 0.001 | 0.59 | 0 | 0.41 | 1.09E-05 |
| 0.001 | 0.95 | 0 | 0.05 | 2.05E-05 |
| 0.001 | 0.6 | 0 | 0.4 | 2.11E-05 |
| 0.001 | 0.61 | 0 | 0.39 | 3.99E-05 |
| 0.001 | 0.62 | 0 | 0.38 | 7.41E-05 |
| 0.001 | 0.94 | 0 | 0.06 | 0.000131786 |
| 0.001 | 0.63 | 0 | 0.37 | 0.000134747 |
| 0.001 | 0.64 | 0 | 0.36 | 0.000240022 |
| 0.001 | 0.65 | 0 | 0.35 | 0.00041857 |
| 0.001 | 0.93 | 0 | 0.07 | 0.000571298 |
| 0.001 | 0.66 | 0 | 0.34 | 0.000714296 |
| 0.001 | 0.67 | 0 | 0.33 | 0.001192219 |
| 0.001 | 0.92 | 0 | 0.08 | 0.00184529 |
| 0.001 | 0.68 | 0 | 0.32 | 0.001945128 |
| 0.001 | 0.69 | 0 | 0.31 | 0.003100024 |
| 0.001 | 0.91 | 0 | 0.09 | 0.00474466 |
| 0.001 | 0.7 | 0 | 0.3 | 0.004822566 |
| 0.001 | 0.71 | 0 | 0.29 | 0.007316707 |
| 0.001 | 0.9 | 0 | 0.1 | 0.010165211 |
| 0.001 | 0.72 | 0 | 0.28 | 0.010815739 |
| 0.001 | 0.73 | 0 | 0.27 | 0.015560463 |
| 0.001 | 0.89 | 0 | 0.11 | 0.018752578 |
| 0.001 | 0.74 | 0 | 0.26 | 0.021760735 |
| 0.001 | 0.75 | 0 | 0.25 | 0.029538931 |
| 0.001 | 0.88 | 0 | 0.12 | 0.030523601 |
| 0.001 | 0.76 | 0 | 0.24 | 0.038858572 |
| 0.001 | 0.87 | 0 | 0.13 | 0.044660732 |
| 0.001 | 0.77 | 0 | 0.23 | 0.049448343 |
| 0.001 | 0.86 | 0 | 0.14 | 0.059598733 |
| 0.001 | 0.78 | 0 | 0.22 | 0.060740148 |
| 0.001 | 0.79 | 0 | 0.21 | 0.07184706 |
| 0.001 | 0.85 | 0 | 0.15 | 0.073380296 |
| 0.001 | 0.8 | 0 | 0.2 | 0.081609113 |
| 0.001 | 0.84 | 0 | 0.16 | 0.084138927 |
| 0.001 | 0.81 | 0 | 0.19 | 0.088727309 |
| 0.001 | 0.83 | 0 | 0.17 | 0.090531145 |
| 0.001 | 0.82 | 0 | 0.18 | 0.091985913 |

**Table S9f** Posterior probabilities of rates of clonality, selfing, and outcrossing using changes of genotype frequencies between temporal samples (CEMP) gathered from Bagley Lake on 30 June 2022 and 5 June 2025. Outcrossing rate is calculated as 1 - clonal rate - selfing rate.

| **Mutation rate** | **Rate of clonality** | **Selfing rate** | **Outcrossing rate** | **Posterior probability** |
| --- | --- | --- | --- | --- |
| 0.001 | 0 | 0 | 1 | 8.52E-49 |
| 0.001 | 0.01 | 0 | 0.99 | 1.47E-47 |
| 0.001 | 0.02 | 0 | 0.98 | 2.21E-46 |
| 0.001 | 0.03 | 0 | 0.97 | 2.96E-45 |
| 0.001 | 0.04 | 0 | 0.96 | 3.57E-44 |
| 0.001 | 0.05 | 0 | 0.95 | 3.93E-43 |
| 0.001 | 0.06 | 0 | 0.94 | 3.98E-42 |
| 0.001 | 0.07 | 0 | 0.93 | 3.74E-41 |
| 0.001 | 0.08 | 0 | 0.92 | 3.28E-40 |
| 0.001 | 0.09 | 0 | 0.91 | 2.69E-39 |
| 0.001 | 0.1 | 0 | 0.9 | 2.08E-38 |
| 0.001 | 0.11 | 0 | 0.89 | 1.52E-37 |
| 0.001 | 0.12 | 0 | 0.88 | 1.06E-36 |
| 0.001 | 0.13 | 0 | 0.87 | 7.00E-36 |
| 0.001 | 0.14 | 0 | 0.86 | 4.42E-35 |
| 0.001 | 0.15 | 0 | 0.85 | 2.67E-34 |
| 0.001 | 0.16 | 0 | 0.84 | 1.55E-33 |
| 0.001 | 0.17 | 0 | 0.83 | 8.62E-33 |
| 0.001 | 0.18 | 0 | 0.82 | 4.62E-32 |
| 0.001 | 0.19 | 0 | 0.81 | 2.39E-31 |
| 0.001 | 0.2 | 0 | 0.8 | 1.20E-30 |
| 0.001 | 0.21 | 0 | 0.79 | 5.80E-30 |
| 0.001 | 0.22 | 0 | 0.78 | 2.72E-29 |
| 0.001 | 0.23 | 0 | 0.77 | 1.24E-28 |
| 0.001 | 0.24 | 0 | 0.76 | 5.48E-28 |
| 0.001 | 0.25 | 0 | 0.75 | 2.36E-27 |
| 0.001 | 0.26 | 0 | 0.74 | 9.86E-27 |
| 0.001 | 0.27 | 0 | 0.73 | 4.03E-26 |
| 0.001 | 0.28 | 0 | 0.72 | 1.60E-25 |
| 0.001 | 0.29 | 0 | 0.71 | 6.23E-25 |
| 0.001 | 0.3 | 0 | 0.7 | 2.36E-24 |
| 0.001 | 0.31 | 0 | 0.69 | 8.78E-24 |
| 0.001 | 0.32 | 0 | 0.68 | 3.19E-23 |
| 0.001 | 0.33 | 0 | 0.67 | 1.13E-22 |
| 0.001 | 0.34 | 0 | 0.66 | 3.95E-22 |
| 0.001 | 0.35 | 0 | 0.65 | 1.35E-21 |
| 0.001 | 0.36 | 0 | 0.64 | 4.51E-21 |
| 0.001 | 0.37 | 0 | 0.63 | 1.48E-20 |
| 0.001 | 0.38 | 0 | 0.62 | 4.78E-20 |
| 0.001 | 0.39 | 0 | 0.61 | 1.52E-19 |
| 0.001 | 0.4 | 0 | 0.6 | 4.72E-19 |
| 0.001 | 0.41 | 0 | 0.59 | 1.44E-18 |
| 0.001 | 0.42 | 0 | 0.58 | 4.35E-18 |
| 0.001 | 0.43 | 0 | 0.57 | 1.29E-17 |
| 0.001 | 0.44 | 0 | 0.56 | 3.75E-17 |
| 0.001 | 0.45 | 0 | 0.55 | 1.08E-16 |
| 0.001 | 0.46 | 0 | 0.54 | 3.04E-16 |
| 0.001 | 0.47 | 0 | 0.53 | 8.48E-16 |
| 0.001 | 0.48 | 0 | 0.52 | 2.33E-15 |
| 0.001 | 0.49 | 0 | 0.51 | 6.29E-15 |
| 0.001 | 0.5 | 0 | 0.5 | 1.68E-14 |
| 0.001 | 0.51 | 0 | 0.49 | 4.41E-14 |
| 0.001 | 0.52 | 0 | 0.48 | 1.14E-13 |
| 0.001 | 0.53 | 0 | 0.47 | 2.93E-13 |
| 0.001 | 0.54 | 0 | 0.46 | 7.40E-13 |
| 0.001 | 0.55 | 0 | 0.45 | 1.84E-12 |
| 0.001 | 0.56 | 0 | 0.44 | 4.54E-12 |
| 0.001 | 0.57 | 0 | 0.43 | 1.10E-11 |
| 0.001 | 0.58 | 0 | 0.42 | 2.65E-11 |
| 0.001 | 0.59 | 0 | 0.41 | 6.27E-11 |
| 0.001 | 0.6 | 0 | 0.4 | 1.47E-10 |
| 0.001 | 0.61 | 0 | 0.39 | 3.39E-10 |
| 0.001 | 0.62 | 0 | 0.38 | 7.75E-10 |
| 0.001 | 0.63 | 0 | 0.37 | 1.75E-09 |
| 0.001 | 0.64 | 0 | 0.36 | 3.90E-09 |
| 0.001 | 0.65 | 0 | 0.35 | 8.59E-09 |
| 0.001 | 0.66 | 0 | 0.34 | 1.87E-08 |
| 0.001 | 0.67 | 0 | 0.33 | 4.02E-08 |
| 0.001 | 0.68 | 0 | 0.32 | 8.54E-08 |
| 0.001 | 0.69 | 0 | 0.31 | 1.79E-07 |
| 0.001 | 0.7 | 0 | 0.3 | 3.72E-07 |
| 0.001 | 0.71 | 0 | 0.29 | 7.63E-07 |
| 0.001 | 0.72 | 0 | 0.28 | 1.54E-06 |
| 0.001 | 0.73 | 0 | 0.27 | 3.09E-06 |
| 0.001 | 0.74 | 0 | 0.26 | 6.10E-06 |
| 0.001 | 0.75 | 0 | 0.25 | 1.19E-05 |
| 0.001 | 0.76 | 0 | 0.24 | 2.29E-05 |
| 0.001 | 0.77 | 0 | 0.23 | 4.35E-05 |
| 0.001 | 0.78 | 0 | 0.22 | 8.15E-05 |
| 0.001 | 0.79 | 0 | 0.21 | 0.000150578 |
| 0.001 | 0.8 | 0 | 0.2 | 0.000274207 |
| 0.001 | 0.81 | 0 | 0.19 | 0.000491858 |
| 0.001 | 0.82 | 0 | 0.18 | 0.000868453 |
| 0.001 | 1 | 0 | 0 | 0.00093396 |
| 0.001 | 0.83 | 0 | 0.17 | 0.001508141 |
| 0.001 | 0.84 | 0 | 0.16 | 0.00257336 |
| 0.001 | 0.85 | 0 | 0.15 | 0.004309324 |
| 0.001 | 0.86 | 0 | 0.14 | 0.007072029 |
| 0.001 | 0.87 | 0 | 0.13 | 0.011353769 |
| 0.001 | 0.88 | 0 | 0.12 | 0.017792929 |
| 0.001 | 0.89 | 0 | 0.11 | 0.027143771 |
| 0.001 | 0.99 | 0 | 0.01 | 0.029510162 |
| 0.001 | 0.9 | 0 | 0.1 | 0.040167653 |
| 0.001 | 0.91 | 0 | 0.09 | 0.057393616 |
| 0.001 | 0.92 | 0 | 0.08 | 0.078694942 |
| 0.001 | 0.98 | 0 | 0.02 | 0.080726423 |
| 0.001 | 0.93 | 0 | 0.07 | 0.102662969 |
| 0.001 | 0.97 | 0 | 0.03 | 0.12381523 |
| 0.001 | 0.94 | 0 | 0.06 | 0.12587134 |
| 0.001 | 0.95 | 0 | 0.05 | 0.142368109 |
| 0.001 | 0.96 | 0 | 0.04 | 0.144145187 |

**Table S10** Intervals of inferred rates of clonality, selfing, and outcrossing from the genotype frequencies. Sampling date 1, first date of sampling in comparison; sampling date 2, second date of sampling in comparison; days elapsed, number of days between sampling date 1 and sampling date 2; maximum posterior clonality, the most likely rate of clonality occurring in the population between the two time points, based on observed changes in genotype frequencies; maximum posterior selfing, the most likely rate of selfing occurring in the population between the two time points, based on observed changes in genotype frequencies; maximum posterior outcrossing, the most likely rate of outcrossing occurring in the population between the two time points, based on observed changes in genotype frequencies.

| **Sampling Date 1** | **Sampling Date 2** | **Days Elapsed** | **Maximum posterior clonality** | **Maximum posterior selfing** | **Maximum posterior outcrossing** |
| --- | --- | --- | --- | --- | --- |
| 2 June 2022 | 23 June 2022 | 21 | 0 | 0.97 | 0.03 |
| 2 June 2022 | 30 Jun 2022 | 28 | 0 | 0.95 | 0.05 |
| 2 June 2022 | 5 June 2025 | 1099 | 0.10 | 0.90 | 0 |
| 23 June 2022 | 30 Jun 2022 | 7 | 0 | 0.90 | 0.10 |
| 23 June 2022 | 5 June 2025 | 1078 | 0 | 0.88 | 0.12 |
| 30 Jun 2022 | 5 June 2025 | 1071 | 0 | 0.97 | 0.03 |

**Table S11a** Significance of observed heterozygosity distribution between sampling dates. Significance was determined using a Kruskal-Wallis test, H = 3.4455, p = 4.8621x10^-01^. Sampling date 1, the first sampling period in the comparison; sampling date 2, the second sampling period in the comparison; U, the probability that a random value from the second sampling date will be greater than a random value from the first sampling group; p_uncorrected, the calculated probability that the difference between the two sampling dates would arise by chance; p_bonferroni, the bonferroni correction applied to correct for testing multiple hypotheses; significant, whether or not the Bonferroni corrected p-value is significant (p>0.05).

| Sampling date 1 | Sampling date 2 | U | p_uncorrected | p_bonferroni | Significant |
| --- | --- | --- | --- | --- | --- |
| 2 June 2022 | 23 June 2022 | 36.5 | 0.136881 | 1.0 | FALSE |
| 2 June 2022 | 30 June 2022 | 28.0 | 0.700852 | 1.0 | FALSE |
| 2 June 2022 | 5 June 2025 | 27.0 | 0.798081 | 1.0 | FALSE |
| 2 June 2022 | Overall | 26.0 | 0.901515 | 1.0 | FALSE |
| 23 June 2022 | 30 June 2022 | 15.0 | 0.243837 | 1.0 | FALSE |
| 23 June 2022 | 5 June 2025 | 14.0 | 0.195844 | 1.0 | FALSE |
| 23 June 2022 | Overall | 14.0 | 0.196347 | 1.0 | FALSE |
| 30 June 2022 | 5 June 2025 | 30.0 | 0.521527 | 1.0 | FALSE |
| 30 June 2022 | Overall | 22.0 | 0.797863 | 1.0 | FALSE |
| 5 June 2025 | Overall | 22.0 | 0.608886 | 1.0 | FALSE |

**Table S11b** Significance of expected heterozygosity distribution between sampling dates. Significance was determined using a Kruskal-Wallis test, H = 2.0871, p = 7.1975x10^-01^. Sampling date 1, the first sampling period in the comparison; sampling date 2, the second sampling period in the comparison; U, the probability that a random value from the second sampling date will be greater than a random value from the first sampling group; p_uncorrected, the calculated probability that the difference between the two sampling dates would arise by chance; p_bonferroni, the bonferroni correction applied to correct for testing multiple hypotheses; significant, whether or not the Bonferroni corrected p-value is significant (p>0.05).

| Sampling date 1 | Sampling date 2 | U | p_uncorrected | p_bonferroni | Significant |
| --- | --- | --- | --- | --- | --- |
| 2 June 2022 | 23 June 2022 | 25.0 | 1.000000 | 1.0 | FALSE |
| 2 June 2022 | 30 June 2022 | 34.0 | 0.259324 | 1.0 | FALSE |
| 2 June 2022 | 5 June 2025 | 26.0 | 0.901515 | 1.0 | FALSE |
| 2 June 2022 | Overall | 24.0 | 1.000000 | 1.0 | FALSE |
| 23 June 2022 | 30 June 2022 | 31.0 | 0.455711 | 1.0 | FALSE |
| 23 June 2022 | 5 June 2025 | 24.0 | 1.000000 | 1.0 | FALSE |
| 23 June 2022 | Overall | 23.0 | 0.901515 | 1.0 | FALSE |
| 30 June 2022 | 5 June 2025 | 16.0 | 0.317599 | 1.0 | FALSE |
| 30 June 2022 | Overall | 15.0 | 0.259324 | 1.0 | FALSE |
| 5 June 2025 | Overall | 21.0 | 0.710373 | 1.0 | FALSE |

**Table S11c** Significance of inbreeding coefficient distribution between sampling dates. Significance was determined using a Kruskal-Wallis test, H = 4.6242, p = 3.2808x10^-01^. Sampling date 1, the first sampling period in the comparison; sampling date 2, the second sampling period in the comparison; U, the probability that a random value from the second sampling date will be greater than a random value from the first sampling group; p_uncorrected, the calculated probability that the difference between the two sampling dates would arise by chance; p_bonferroni, the bonferroni correction applied to correct for testing multiple hypotheses; significant, whether or not the Bonferroni corrected p-value is significant (p>0.05).

| **Sampling date 1** | **Sampling date 2** | **U** | **p_uncorrected** | **p_bonferroni** | **Significant** |
| --- | --- | --- | --- | --- | --- |
| 2 June 2022 | 23 June 2022 | 13.0 | 0.155249 | 1.0 | FALSE |
| 2 June 2022 | 30 June 2022 | 22.0 | 0.804779 | 1.0 | FALSE |
| 2 June 2022 | 5 June 2025 | 18.0 | 0.455711 | 1.0 | FALSE |
| 2 June 2022 | Overall | 20.0 | 0.620047 | 1.0 | FALSE |
| 23 June 2022 | 30 June 2022 | 37.0 | 0.121037 | 1.0 | FALSE |
| 23 June 2022 | 5 June 2025 | 36.0 | 0.155249 | 1.0 | FALSE |
| 23 June 2022 | Overall | 37.0 | 0.121037 | 1.0 | FALSE |
| 30 June 2022 | 5 June 2025 | 18.0 | 0.455711 | 1.0 | FALSE |
| 30 June 2022 | Overall | 21.0 | 0.710373 | 1.0 | FALSE |
| 5 June 2025 | Overall | 27.0 | 0.804779 | 1.0 | FALSE |

**Table S12** Temporal *F_ST_* between sampling dates. Sampling date 1, the first date in the comparison; sampling date 2, the second date in the comparison; Temporal *F_ST_*, the *F_ST_* values calculated between those dates.

|  | **Sampling Date 1** | **Sampling Date 2** | **Temporal *F_ST_*** |
| --- | --- | --- | --- |
|  | 2 June 2022 | 23 June 2022 | 0.0282 |
| **June 2022** | 2 June 2022 | 30 June 2022 | 0.0454 |
|  | 23 June 2022 | 30 June 2022 | 0.0814 |
|  | 2 June 2022 | 5 June 2025 | 0.0176 |
| **Between 2022 and 2025** | 23 June 2022 | 5 June 2025 | 0.0347 |
|  | 30 June 2022 | 5 June 2025 | 0.0404 |

**Table S13** Significance of pairwise linkage distribution (LD) between sampling dates. Significance was determined using a Kruskal-Wallis test, H = 71.7118, p = 9.8740x10-15. Sampling date 1, the first sampling period in the comparison; sampling date 2, the second sampling period in the comparison; U, the probability that a random value from the second sampling date will be greater than a random value from the first sampling group; p_uncorrected, the calculated probability that the difference between the two sampling dates would arise by chance; p_bonferroni, the bonferroni correction applied to correct for testing multiple hypotheses; significant, whether or not the Bonferroni corrected p-value is significant (p>0.05).

| **Sampling date 1** | **Sampling date 2** | **U** | **p_uncorrected** | **p_bonferroni** | **Significant** |
| --- | --- | --- | --- | --- | --- |
| 30 June 2022 | Overall | 432.0 | 1.109191e-07 | 0.000001 | TRUE |
| 5 June 2025 | Overall | 422.0 | 4.274603e-07 | 0.000004 | TRUE |
| 23 June 2022 | Overall | 421.0 | 4.875515e-07 | 0.000005 | TRUE |
| 2 June 2022 | 23 June 2022 | 21.0 | 5.557498e-07 | 0.000006 | TRUE |
| 23 June 2022 | 5 June 2025 | 418.0 | 7.207746e-07 | 0.000007 | TRUE |
| 2 June 2022 | 30 June 2022 | 37 | 4.154483e-06 | 0.000042 | TRUE |
| 2 June 2022 | Overall | 393.0 | 1.512999e-05 | 0.000151 | TRUE |
| 23 June 2022 | 30 June 2022 | 361.0 | 4.286215e-04 | 0.004286 | TRUE |
| 30 June 2022 | 5 June 2025 | 355.0 | 7.493143e-04 | 0.007493 | TRUE |
| 2 June 2022 | 5 June 2025 | 149.0 | 7.408994e-02 | 0.740899 | FALSE |
